# Sensitive, high-throughput HLA-I and HLA-II immunopeptidomics using parallel accumulation-serial fragmentation mass spectrometry

**DOI:** 10.1101/2023.03.10.532106

**Authors:** Kshiti Meera Phulphagar, Claudia Ctortecka, Alvaro Sebastian Vaca Jacome, Susan Klaeger, Eva K. Verzani, Gabrielle M. Hernandez, Namrata Udeshi, Karl Clauser, Jennifer Abelin, Steven A Carr

## Abstract

Comprehensive, in-depth identification of the human leukocyte antigen HLA-I and HLA-II tumor immunopeptidome can inform the development of cancer immunotherapies. Mass spectrometry (MS) is powerful technology for direct identification of HLA peptides from patient derived tumor samples or cell lines. However, achieving sufficient coverage to detect rare, clinically relevant antigens requires highly sensitive MS-based acquisition methods and large amounts of sample. While immunopeptidome depth can be increased by off-line fractionation prior to MS, its use is impractical when analyzing limited amounts of primary tissue biopsies. To address this challenge, we developed and applied a high throughput, sensitive, single-shot MS-based immunopeptidomics workflow that leverages trapped ion mobility time-of-flight mass spectrometry on the Bruker timsTOF SCP. We demonstrate >2-fold improved coverage of HLA immunopeptidomes relative to prior methods with up to 15,000 distinct HLA-I and HLA-II peptides from 4e7 cells. Our optimized single-shot MS acquisition method on the timsTOF SCP maintains high coverage, eliminates the need for off-line fractionation and reduces input requirements to as few as 1e6 A375 cells for > 800 distinct HLA-I peptides. This depth is sufficient to identify HLA-I peptides derived from cancer-testis antigen, and novel/unannotated open reading frames. We also apply our optimized single-shot SCP acquisition methods to tumor derived samples, enabling sensitive, high throughput and reproducible immunopeptidome profiling with detection of clinically relevant peptides from less than 4e7 cells or 15 mg wet weight tissue.

## Introduction

Anti-tumor CD8+ and CD4+ T cells can identify tumor cells through the recognition of peptide antigens presented by human leukocyte antigen class I (HLA-I) and class II (HLA-II) cell surface molecules, respectively. Immunotherapies that harness the ability of T cells to recognize aberrant cells in the body have shown great clinical potential (1). Thus, personalized cancer vaccines and cell therapies targeting HLA peptides that contain patient specific somatic mutations (neoantigens) and tumor antigens have shown success in early-stage clinical trials (2–4). Additionally, tumor specific noncanonical antigens, derived from translation of non-protein-coding regions, or from noncanonical transcription and/or translation could have great potential as novel therapeutic targets (5–7). Therefore, it is crucial to have technologies that enable comprehensive analysis of peptides presented by both HLA-I and HLA-II molecules on tumors.

The highly polymorphic nature of HLA alleles makes their peptide binding rules complex to characterize. Almost all cells in the body express HLA-I molecules that present peptides of length 8-12mer with specific amino acids, usually at position 2 and the termini of each peptide, called anchor residues that interact with the HLA-I binding groove (8). In contrast, HLA-II molecules present longer peptides, usually 12-25 amino acids in length. HLA-II peptides have allele-specific binding registers that usually contain four anchor residues that can be anywhere within the peptide sequence and are generally expressed by antigen presenting cells or epithelial cells exposed to interferon gamma (8). Due to these expression patterns and the well documented cytotoxic activity of CD8+ T cells, HLA-I tumor immunopeptidomes are most frequently studied (1, 5, 8). Nonetheless, both HLA-I and HLA-II tumor antigen profiling are needed, as the latter play an important role in anti-tumor immunity in both HLA-II and non-HLA-II expressing tumors (9–11).

MS has proven to be a powerful technology to define and characterize HLA-I and HLA-II immunopeptidomes. In 1992 Hunt et al. sequenced eight HLA-I peptides from 2 billion cells post fractionation using microcapillary high-performance liquid chromatography-electrospray ionization-tandem MS (12). Current methods and instrumentation for MS based immunopeptidome analyses now enables 1000’s of HLA-I and HLA-II peptides to be identified from similar starting amounts of material (13–15). Yet achieving significant depth (i.e., >10,000 HLA peptides) to identify rare, clinically relevant antigens by LC-MS/MS-based immunopeptidomics from a single HLA immunopurification of <500 million cells remains a challenge (16–18). More recently, microscaled basic reversed phase (brp) fractionation of enriched HLA peptides and ion mobility (IM) coupled to LC–MS/MS enabled deep coverage of the HLA peptidome and detection of neoantigen peptides from only 100 million cells or 150 mg wet tumor weight (19). However, reducing sample complexity through offline-fractionation increases the number of LC-MS/MS runs per sample 3-6-fold and is not suitable for sample amounts lower than 100 mg due to accompanying adsorptive peptide losses (19). Current methods therefore are not sufficiently scalable for immunopeptidome profiling of large sample cohorts or primary patient samples where tissue amounts are limited.

With the goal to develop MS-based immunopeptidomics methods with throughput sufficient to analyze hundreds of patient samples while achieving high depth, we investigated single-shot LC-MS/MS complemented by IM separation. Here we comparatively evaluate single-shot methods on the Thermo Scientific Orbitrap Exploris 480 with FAIMS Pro (Exploris + FAIMS) and the Bruker timsTOF SCP (SCP) mass spectrometers. Acquisition methods for the SCP were optimized and evaluated for the analysis of both HLA-I and HLA-II peptides capitalizing on trapped ion-mobility mass spectrometry (TIMS) separation and parallel accumulation-serial fragmentation (PASEF) (20–22). We found that use of the SCP, a dedicated low-input instrument, improved depth of HLA-I and HLA-II immunopeptidomes >2 fold while increasing throughput nearly 4-fold relative to offline brp fractionation. Our improved workflow demonstrates useful immunopeptidome coverage from only 1e6 cells or 15 mg tumor tissue and up to 15,000 distinct HLA-I and HLA-II peptides from 4e7 cells. These optimized single-shot immunopeptidomics methods developed using the SCP instrument have sufficient throughput and reproducibility for profiling the immunopeptidome at scale, thus enabling the analysis of large clinical sample cohorts for the detection of clinically relevant antigens.

### Experimental Procedures

#### Culture of A375 melanoma cells and 2D pancreatic ductal carcinoma derived from primary human samples

Conditions for growth and *in vitro* propagation of A375 melanoma were as per ATCC guidelines and for pancreatic ductal adenocarcinoma tumor cell (PDAC) lines were described previously (23). For all cell lines, cells were counted by automated cell counting after trypan blue staining, specified cell amounts ranging from 0.5 to 100 million cells or 1 billion cells for bulk HLA peptide enrichment were harvested by trypsinization (Trypsin-EDTA 0.25%, Gibco™ 25200056), pelleted and rinsed in PBS twice. Pellets were either snap frozen or used directly for HLA enrichment.

#### Collection of melanoma tumor samples

Melanoma tumor samples were collected as part of the NIH/NCI CPTAC consortium (https://proteomics.cancer.gov/programs/cptac) with protocols mandated by the CPTAC program office. Data collection and analysis in this study was performed in accordance with the Declaration of Helsinki and Institutional review boards at tissue source sites reviewed protocols and consent documentation adhering to the CPTAC guidelines. Information on participant compensation is not available to the investigators.

#### HLA-I Peptide Enrichment and Peptide Elution

Cell pellets were lysed in lysis buffer containing 20 mM Tris, pH 8.0, 100 mM NaCl, 6 mM MgCl_2_, 1 mM EDTA, 60 mM Octyl β-d-glucopyranoside, 0.2 mM Iodoacetamide, 1.5% Triton X-100, 1x Complete Protease Inhibitor Tablet-EDTA free, 1 mM PMSF in total of 1.2 ml lysate per 50 million cells as described previously (24). Each lysate was transferred to an Eppendorf tube, incubated on ice for 30 min with 2 µL of Benzonase (Thomas Scientific, E1014-25KU) and inverted every 5 min. Then lysates were centrifuged at 15,000 rcf for 20 min at 4°C and the supernatants were transferred to 96 deep-well plates (Cytiva, 7701-5200).

Immunoprecipitation of HLA-I only or both HLA-I and II was performed sequentially as follows: ∼37.5 µL pre-washed gamma bind sepharose beads (Millipore Sigma, GE17-0886-01) and 15 µL of HLA-II antibody mix (9 µL TAL-1B5 (Abcam, ab20181), 3 µL EPR11226 (Abcam, ab157210), 3 µL B-K27 (Abcam, ab47342) were added to lysates in each well and plates were sealed with 96-well square sealing mats (Thermo-Fisher Scientific, AB0675). HLA complexes were captured on the beads by incubating on a rotor at 4°C for 3 hr. Following the incubation beads and lysates were transferred to a pre-washed 10 μm PE fritted plate (Agilent, S7898A) stacked on top of a fresh 96 deep-well plate. Beads with HLA-II peptides complexes were retained on the filter plate while lysates were collected in the fresh 96-well plate. For HLA-I enrichment, 37.5 µL pre-washed beads and 15 µL of HLA-I antibody (W6/32) (Abcam 22432 or Novus Biologicals NB100-64775) were added to the filtered lysates in each well of the 96-well plate and incubated at 4°C for 3hr.

Beads were transferred and washed on a 10 μm PE fritted plate placed on a positive pressure manifold, as described previously (24). HLA peptides were eluted and desalted from beads using the tC18 40 mg Sep-Pak desalting plate (Waters, Milford, MA) as recommended by the manufacturer. The peptides were eluted from the Sep-Pak desalt plate using 250 µL 15% ACN/1% FA and 2x 250 µL of 50% ACN/1% FA in 96 deep-well plates, snap frozen, and lyophilized.

Dried peptides were stored at -80°C until micro-scaled brp separation on SDB-XC stage tips as previously described (19). Briefly, peptides were reconstituted in 200 µl 3% ACN/5% FA, loaded on equilibrated Stage-tips with 2 punches SDB-XC material (CDS analytical, previously Empore 3M, 13-110-059). After 3 washes with 1% FA, HLA-I and HLA-II peptides were eluted with increasing ACN concentrations of 5/10/30% in 0.1% NH4OH at pH 10, to 96-well PCR plates (60180-P210B), frozen and dried down by vacuum centrifugation.

#### LC-MS/MS analysis

Peptides were reconstituted in 3% ACN/5% FA prior to loading onto an analytical column (35 cm, 1.9µm C18 (Dr. Maisch HPLC GmbH), packed in-house PicoFrit 75 µm inner diameter, 10 µm emitter (New Objective)). For acquisition on the Exploris + FAIMS, peptides were eluted with a linear gradient (EasyNanoLC 1200, Thermo Fisher Scientific) ranging from 6-30% Solvent B (0.1% FA/90% ACN) over 84 min, 30-90% B over 9 min and held at 90% B for 5 min at 200 nL/min. MS1 spectra were acquired from 350-1700 m/z at 60,000 resolution and collected until either 100% normalized automatic gain control (AGC) target or a maximum injection time of 50 ms was reached. The RF amplitude applied to the RF lens was set to 40% and advanced peak determination was enabled. Monoisotopic peak detection was set to “peptide,” and “relax restrictions” were enabled. The precursor fit threshold was set to 50% at 1.2 m/z fit window and dynamic exclusion was 10 sec at 10 ppm. Potential precursors above 1e4 were isolated in data dependent acquisition (DDA) mode with a 1.1 m/z window, until 50% AGC target or 120 ms maximum injection time was reached. HLA-I precursors were fragmented at 30% normalized higher energy collisional dissociation (HCD) and HLA-II at 34%, both with 15,000 resolution. When the FAIMS interface was used, spray voltage was increased to 1900 V at “standard resolution”. FAIMS compensation voltages (CVs) of -50 and -70, each with a cycle time of 1.5 s were used. Instrument performance was evaluated using an in-house 10 ng jurkat proteome digest (protein quantification), and data were acquired above a peptide yield threshold of >16,000 peptides for a 120 min LC gradient.

For acquisition on the SCP, peptides were separated with a linear, stepped gradient with the Bruker nanoElute ranging from 2-15 Solvent B (0.1% FA in ACN) over 60 minutes, 15-23% in 30 minutes, 23-35% in 10 minutes, 35-80% in 10 minutes and held at 80% for 10 minutes at 400 nL/min. MS1 scans were acquired from 100-1700 m/z and 1/K0 = 1.7 Vs cm^-2^ to 0.6 Vs cm^-2^ for HLA-I or 1/K0 = 1.3 Vs cm^-2^ to 0.6 Vs cm^-2^ for HLA-II in DDA-PASEF mode. Ten PASEF ramps were acquired with an accumulation and ramp time of 166 ms or as stated otherwise. Precursor above the minimum intensity threshold of 1000 were isolated with 2 Th at < 700 m/z or 3 Th >800 m/z and re-sequenced until a target intensity of 10,000 considering a dynamic exclusion of 40 s or as stated otherwise. The collision energy (CE) was lowered linearly as a function of increasing mobility starting from 55 eV at 1/K0 = 1.6 Vs cm^-2^ to 10 eV at 1/K0 = 0.6 Vs cm^-2^ or as specified. Standard polygon placement was adapted for singly charged HLA-I peptide species and as detailed in the result section below. Instrument performance was evaluated using an in-house 50 ng jurkat proteome digest (protein quantification), and data were acquired above a peptide yield threshold of >85,000 peptides for a 120 min LC gradient.

#### HLA Peptide Search and Identification

Mass spectra were interpreted using the Spectrum Mill (SM) software package, version 8.01 (Broad Institute; proteomics.broadinstitute.org) as previously described (19, 25, 26) with modifications detailed below. Briefly, only MS/MS spectra with precursor sequence MH+ in the range 700-2000 for HLA-I and 700-4000 for HLA-II, a precursor charge 1-4 for HLA-I and 2-6 for HLA-II, or a minimum of <5 detected peaks were extracted. Merging of similar spectra with the same precursor m/z acquired in the same chromatographic peak was enabled. For Exploris merging was limited to spectra with precursor selection purity of greater than 75%. Precursor selection purity calculates the proportion of ion current in the isolation window of a high resolution MS1 scan represented by the isotope cluster of precursor ions assigned to the resulting MS/MS scan. MS/MS spectra with a sequence tag length >1 (i.e., minimum of three masses separated by the in-chain masses of two amino acids) were searched with no-enzyme specificity; instrument: ESI-QEXACTIVEHCD-HLA-v3; fixed modifications: carbamidomethylation of cysteine, variable modifications: oxidation of methionine, pyroglutamic acid at peptide N-terminal glutamine, cysteinylation, protein N-terminal acetylation and deamidation; precursor mass tolerance of±10 ppm; product mass tolerance of ±10 ppm for Exploris or ±15 ppm for SCP data; minimum matched peak intensity (percent scored peak intensity or %SPI) of 40%. Scored percent intensity (SPI) is the percent of product ion intensity (after peak detection) that is matched to a scored ion type.

MS/MS spectra were searched against the human reference proteome Gencode 34 (HLA-I) or 42 (HLA-II) (ftp.ebi.ac.uk/pub/databases/gencode/Gencode_human/release_34/42) with 47,429 or 50,872 non redundant protein coding transcript biotypes mapped to the human reference genome GRCh38, 602 common laboratory contaminants, 2043 curated smORFs (lncRNA and uORFs), 237,427 novel unannotated ORFs (nuORFs) supported by ribosomal profiling nuORF DB v1.037 for a total of 287,501 or 290,944 entries, respectively (6).

Peptide spectrum matches (PSMs) within <1% false discovery rate (% FDR) using the target decoy estimation of the SM autovalidation module were filtered for a sequence length of 8-12 aa or 12-40, a minimum backbone cleavage score (BCS) of 5 or 7 for HLA-I or HLA-II peptides, respectively. BCS is a peptide sequence coverage metric to enforce a uniformly higher minimum sequence coverage for each PSM. The BCS score is a sum after assigning a 1 or 0 between each pair of adjacent amino acids in the sequence (maximum score is peptide length -1) considering all selected ion types to decrease false positive spectra having fragmentation in a limited portion of the peptide by multiple ion types.

PSMs were consolidated to peptides using the SM protein/peptide summary modules case sensitive peptide-distinct mode. A distinct peptide was the single highest scoring PSM of a peptide detected for each sample. Different modification states observed for a peptide were each reported when containing amino acids configured to allow variable modifications; a lowercase letter indicates the variable modification (C-cysteinylated, c-carbamidomethylated). Additionally, precursor fragmentation was evaluated through the percent dissociated intensity (PDI). The PDI reports the intensity of residual precursor and its neutral losses of water and ammonia subtracted from the total peak intensity in the MS/MS spectrum divided by the total peak intensity.

For MS1 quantification of both SCP and Exploris + FAIMS, we used MSFragger 3.6 (27, 28) within FragPipe 19.0 and IonQuant 1.8.9 (29) and Philosopher 4.7.0 (30) to search spectra against the Ensembl v100 (March 2020) database including 71,704 non-redundant protein coding transcript biotypes mapped to the human reference genome GRCh38, 602 common laboratory contaminants, 2043 curated smORFs (lncRNA and uORFs), 237,427 novel unannotated ORFs (nuORFs) supported by ribosomal profiling nuORF DB v1.553 for a total of 311,776 entries and 50% reverse peptide sequences. For this we used a precursors mass tolerance of ±20 ppm for the SCP or ±15 ppm for the Exploris and a fragment mass tolerance of 10 ppm including minimum 5 matched fragment ions. We performed nonspecific enzymatic protein digestion within 7-14 aa per peptide in a mass range of 600-3000 and a precursor charge 1-4. A maximum of 3 variable modifications per peptides were allowed including oxidation of methionine, N-terminal acetylation and deamidation, cysteinylation, pyroglutamic acid at peptide N-terminal glutamine and fixed cysteine carbamidomethylation. For IonQuant we used 10 ppm mass tolerance, 0.05 IM tolerance, 0.4 RT tolerance at a 1% ion, peptide, and protein FDR without match between runs. Data was filtered for 1% FDR on PSM and peptide level and subsequently filtered for peptides common to SM peptideExport.

Identified peptides were filtered for common contaminants, peptides for which both the preceding and C-terminal amino acids were tryptic residues and peptides observed in negative control runs as described previously (25, 26). Subsequent data analysis was performed in the R computational environment.

#### Subset-specific FDR filtering for nuORFs

While the aggregate FDR for was set to < 1%, as described above, FDR for subset of nuORFs (<5% of total of HLA-I peptides), required more stringent score thresholding to reach a suitable subset specific FDR < 1.0%. To this end, we devised and applied subset-specific filtering approaches. Subsets of nuORF types were thresholded independently in the HLA datasets using a 2-step approach. First, PSM scoring metric thresholds were tightened in a fixed manner for all nuORF PSMs so that nuORF distributions for each metric improved to meet or exceed the aggregate distributions. For all ‘omes the fixed thresholds were: minimum score: 7, minimum %SPI: 50%, precursor mass error: +/-5 ppm, minimum backbone cleavage score (BCS): 5, sequence length: 8-12 (HLA-I), 9-50 (HLA-II). Second, individual nuORF type subsets with FDR estimates remaining above 1% were further subject to a grid search to determine the lowest values of BCS (sequence coverage metric) and score (fragment ion assignment metric) that improved FDR to < 1% for each ORF type in the dataset.

#### HLA Peptide Prediction

HLA peptide prediction was performed using HLAthena (hlathena.tools) (26). Unless otherwise specified,peptides were assigned to an allele using a percentile rank cutoff of ≤0.5.

#### Experimental Design and Statistical Rationale

Sample preparation and data acquisition parameters for both instruments are detailed in the sections above. HLA peptides were enriched in bulk from 1e9 A375 cells and aliquots corresponding to 1e7 cells were injected in technical duplicate for method optimization on the SCP (Figure 1; n=2). Titration experiments using 1e6 to 4e7 cell aliquots of the bulk purified HLA-I peptides were analyzed in technical triplicate (Figure 2 & 3; n=3). Direct IP experiments were carried out by enriching HLA peptides from starting amounts of cells corresponding to 1e6-4e7 million cells in triplicates (Figure 4; n=3). ∼15mg wet weight of primary melanoma tumors were used for HLA-I peptide enrichment from five patients in single-shot injections due to tissue constraints (Figure 5; n=5). Sample size was n = 1 for PDAC line and primary melanoma tumor samples per instrument setup due to sample input limitations. Median peptide counts across replicates ± standard deviation and pearson correlation of peptide intensities are shown.

**Figure 1:**
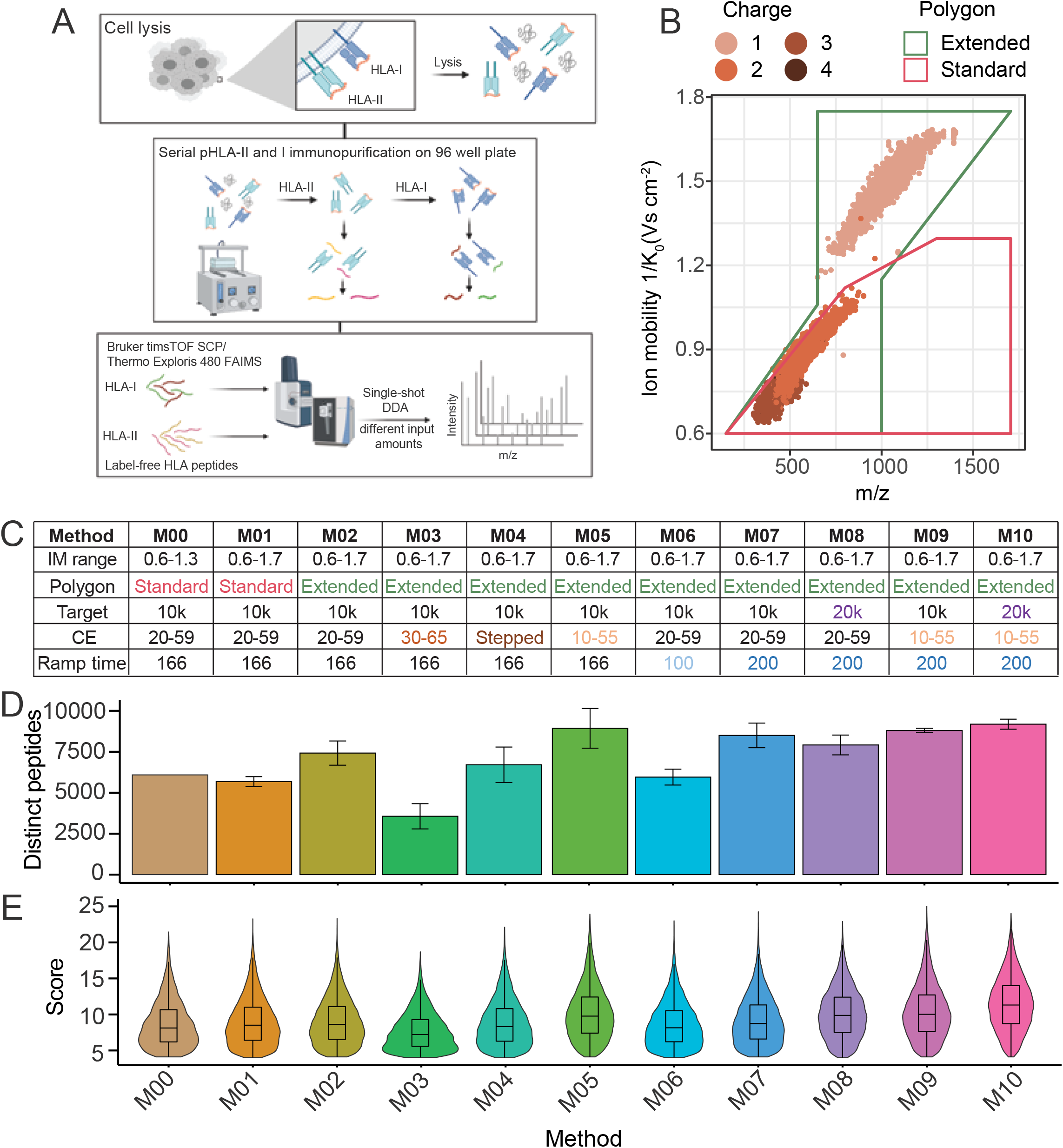
HLA sample preparation overview and timsTOF SCP acquisition method optimization for analysis of HLA-I peptides enables inclusion of singly charged peptides. A, Schematic overview of HLA sample preparation and MS acquisition schemes. Serial HLA-I and HLA-II enrichment, acid elution of peptides from HLA complexes and peptide desalting performed in a semi-automated 96-well format, followed by single-shot DDA analysis of purified HLA-I and HLA-II peptides using our standard workflow with the Exploris + FAIMS or the timsTOF SCP. B, HLA-I PSMs identified on the timsTOF SCP using the standard precursor filter (‘polygon’) in pink andthe extended polygon in green across m/z and IM dimensions. Colors indicate precursor charge state. C, timsTOF SCP parameter overview including, IM range, polygon placement, target intensity value(‘Target’), CE slope, accumulation and ramp time for M00-M10. D, Number of unique HLA-I peptides from 1e7 cell equivalents post-filtering in duplicates with M00-M10 on the timsTOF SCP. Median and standard deviation are shown. E, Corresponding score distributions of HLA-I peptides for methods indicated in D. IM is ion mobility, CE is collisional energy.

**Figure 2:**
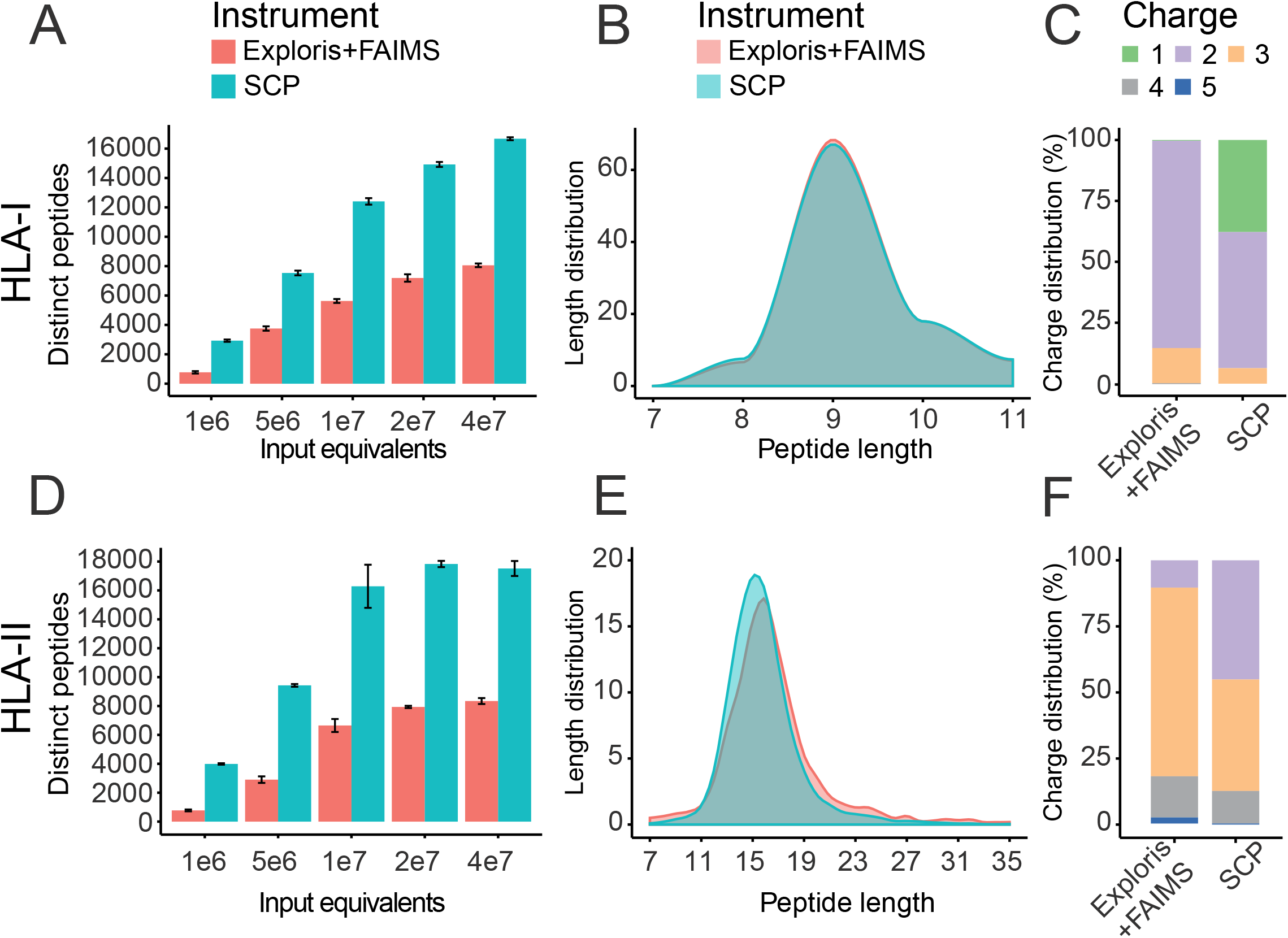
Single-shot acquisition of HLA-I and HLA-II peptides on the timsTOF SCP increases identifications >2-fold compared to Exploris + FAIMS. A, Unique HLA-I peptides identified by single injections on Exploris + FAIMS (red) and timsTOF SCP (blue) from bulk digest at indicated input equivalents. Mean and standard deviation is shown. B, Peptide length distribution across HLA-I peptides identified on the Exploris + FAIMS (red) and the timsTOF SCP (blue) from bulk digest at indicated input equivalents. C, HLA-I peptide charge states for Exploris + FAIMS or timsTOF SCP. D, Unique HLA-II peptides identified by single injections on Exploris + FAIMS (red) and timsTOF SCP and indicated input equivalents (blue). Mean and standard deviation is shown. E, HLA-II peptide length distribution for Exploris + FAIMS (red) and the timsTOF SCP (blue). F, Charge state distribution for HLA-II peptides on Exploris + FAIMS or timsTOF SCP.

## Results

### Single shot HLA-I and HLA-II method optimization and benchmarking

To improve depth, sensitivity, and throughput of our existing immunopeptidomics analysis workflow we focused on optimizing MS acquisition and directly compared single-shot LC-MS/MS DDA acquisition on the SCP to a recently published method using the Exploris + FAIMS (19, 24). We aimed for a single-shot LC-MS/MS analysis to minimize sample handling related to biochemical fractionation and to enable high-throughput analyses of input limited samples. For our initial evaluation, we used 1e7 cell equivalents of HLA-I and HLA-II peptides derived from a bulk preparation of 1e9 A375 cells, a multi-allelic melanoma cell line (Figure 1A). A375 expresses lower levels of TAP genes and has moderate HLA-I and HLA-II expression, making it well suited for optimizing serial HLA immunopeptidome analysis (31). To achieve maximum immunopeptidome depth at high throughput we investigated a wide range of precursor selection, accumulation time and fragmentation parameters to identify the optimal combination for HLA alleles expressed by A375 (Figure 1B, C, Table S1).

In the dual-TIMS analyzer within the SCP, ions are separated by their collisional cross-section (CCS), based on their size, shape, and charge. This allows filtering of the selection potential precursors for MS/MS based on their IM and m/z (32, 33). The ‘standard’ polygon-shaped filter shown in Figure 1B (pink box) is designed to isolate multiply charged precursors and exclude singly charged contaminants in methods optimized for the analysis of tryptic peptides. Tryptic and HLA-II peptides are more similar in length and charge properties than HLA-I peptides are to tryptic peptides. In contrast, HLA-I peptides are shorter, generally 8-12 mers, and allele specific binding motifs that frequently lack basic amino acids, resulting in singly charged precursors with atypical CCS and IM characteristics. To better capture HLA-I peptides, we first expanded the IM range from standard 1/K0 1.3 Vs cm^-2^ to 1/K0 = 1.7 Vs cm^-2^ (M00-M01; Figure 1C). The extension of the ‘standard’ tryptic polygon resulted in comparable numbers of HLA-I peptides identified relative to M00 (Figure 1D-E). We therefore modified the polygon placement to include singly charged precursors with m/z >600 (Figure 1B, green box and Figure 1C “extended polygon” in M02-M10) and observed a 20% increase in peptide IDs (Figure 1D, S1A-B). Upon inclusion of singly charged precursors using the extended polygon within M02, we observed that these precursors had a lower PDI distribution compared to higher charged ones (Figure S2B). As CE on the timsTOF is ramped linearly as a function of IM, we speculated that previously optimized CEs for tryptic peptides are not directly transferable to HLA-I peptides with atypical CCSs, particularly singly charged ones. We tested whether increasing precursor fragmentation could improve identification and linearly increased CEs (30 eV at 1/K0 = 0.6 Vs cm^-2^ to 65 eV at 1/K0 = 1.6 Vs cm^-2^) relative to M02 (M02 vs M03; Figure 1C, S2A). As expected, this slightly increased median PDI by 2%, 4% and 2% for +1, +2 and +3 charged precursors, respectively, but surprisingly this change lowered peptide identifications by over 50% (M02 vs M03; Figure 1E-F, S1A-B, S2B). Therefore, we instead empirically optimized CEs for individual charge states separated by their IM. To accomplish this, we introduced a step to the CE slope, starting with lower CEs at lower IM compared to M02 (1/K0 0.6-1.1 Vs cm^-2^) and a higher CE slope at high IM of method M03 (i.e. 1/K0 1.1-1.6 Vs cm^-2^ for singly charged precursors; Figure S2A). These changes did not result in the expected peptide ID increase, but to the contrary, decreased IDs by 9.5% (M02 vs M04; Figure 1D, S1A-B). We then tested lowering the CE slope (10 eV at 1/K0 = 0.6 Vs cm^-2^ to 55 eV at 1/K0 = 1.6 Vs cm^-2^). This change decreased median PDI by 5%, 34% and 46% for +1, +2 and +3 charged precursors, respectively (M02 vs M05; Figure S1B, E,), but unexpectedly peptide IDs increased by 16% and the % SPI increased by 5% (M02 vs M05; Figure 1C, Figure S1A-C).

Next, to increase scanning speed and thus decrease cycle time, we reduced the standard TIMS ramp time from 166 ms to 100 ms (M02 vs M06; Figure 1C). The TIMS ramp time defines the initial accumulation time of ions in the first TIMS funnel and the subsequent elution time from the second TIMS to the quadrupole for precursor isolation. However, this change decreased peptide identifications by 15% (Figures 1D-E, S1A-B). We next evaluated the effect of increasing the ramp time to 200 ms. Increasing the ion accumulation increases the peptide ions available for fragmentation and thus improves HLA-I peptide identifications by 14% (M02 vs M07; Figure 1C-E, S1A-B). Doubling the target intensity from 10,000 to 20,000 improved the median peptide score by 19% and marginally increased peptide identifications by 5%, presumably by improving the coverage of complementary b and y type ions (M02 vs M08; Figure 1C-E, S1A-E). Based on the collective results obtained from our testing, we designed method M10 that combines optimized polygon filter placement, lower CE slope of 10 eV at 1/K0 = 0.6 Vs cm^-2^ to 55 eV at 1/K0 = 1.6 Vs cm^-2^, 200 ms ramp time and a higher target intensity threshold of 20,000 (Figure 1C-E, Figure S1A-E). M10 yielded the highest number of IDs and median scores, and we therefore considered parameter set M10 as optimal for HLA-I immunopeptidomics of A375 on the SCP.

### Improved coverage of HLA-I and HLA-II immunopeptidomes on the SCP

With the optimized settings described above, we next compared single-shot analyses on the SCP to the Exploris + FAIMS). For this, HLA-I and HLA-II peptides were bulk enriched from 1e9 A375 cells and equivalents of 1e6, 5e6, 1e7, 2e7 and 4e7 cells were injected in triplicates on both instruments (Tables S2, 3, 5 and 6). For HLA-I peptides, we observed a 2-fold increase in identifications using the SCP in comparison to the Exploris + FAIMS at 5e6 cells and higher (Figure 2A). At 4e7 input level, we observed over 17,000 HLA-I peptides on the SCP vs ca. 8000 on the Exploris + FAIMS. At 1e6 cells the optimized SCP method yielded ∼3,000 HLA peptides in comparison to ∼800 peptides on the Exploris + FAIMS, a differential of nearly 4-fold (Figure 2A). The increase in peptide IDs corresponded to a 1.5 fold increase in the number of source proteins represented in the HLA-I immunopeptidome (Figure S3A). Based on their length or charge distributions (Figure 2B-C) and presence of expected anchor residues for A375 alleles (Figure S6D), we confirmed that the additional peptides identified by the SCP are indeed HLA-I bound. Parameter set M10 on the SCP yielded 92%, 70% and 64% median PDI for +1, +2 and +3 charged precursors, respectively. While the median PDI using the SCP was lower by 28% and 35% for +2 and +3 charged precursors than on the Exploris + FAIMS, the scores and sequence coverage were comparable (Figure S4A-C). As expected, more singly charged HLA-I peptides were observed using the SCP than the Exploris + FAIMS as those are generally excluded by the FAIMS CVs used (Figure 2C). Interestingly, spectra on the Exploris + FAIMS are composed predominantly of y type and internal ions, while the SCP yielded more complementary b and y type ion pairs, with few internal ions (Figure S6C). Thus, the SCP enables single shot HLA-I immunopeptidome profiling using input amounts of 1e6 cells and higher due to the 2-fold higher peptide IDs at comparable scores and more complementary b and y type ion pairs compared to the Exploris + FAIMS (Figure 2A, Figure S3C).

As HLA-II peptides are longer than HLA-I peptides and present with higher charge states more similar to tryptic peptides (i.e. +2 to +5), we used a previously published method based on default tryptic acquisition methods on the SCP with standard polygon filter, CE 20 eV at 1/K0 = 0.6 Vs cm-2 to 59 eV at 1/K0 = 1.6 Vs cm^-2^ and target intensity threshold of 20,000 (21). We observed a 2-3 fold increase of HLA-II peptide identifications on the SCP compared to the Exploris + FAIMS (Figure 2D). HLA-II peptide length (12-25 mers) and charge distribution +2 to +5 was observed on the SCP as expected, with a slight shift in the median length (15 vs. 16, Figure 2E-F). Moreover, we observed more peptides with charge state +2 on the SCP, while peptides identified on the Exploris + FAIMS are predominantly of higher charge (i.e. +3 to +5, Figure 2F). We hypothesize that the Exploris + FAIMS is identifying longer and more highly charged peptides because it leverages a charge state specific CE that is higher and likely results in improved fragmentation for longer and high charged peptides. While overall number of HLA-II peptide identifications is 2-fold higher on the SCP, HLA-II peptides on both instruments show comparable scores and sequence coverage (indicated by BCS) to those observed on the Exploris + FAIMS, with slightly lower precursor dissociation of 24%, 7%, 3% and 2% for +2, +3, +4 and +5 precursors, respectively (Figure S5A-B). In summary, across input of 1e6 to 4e7 A375 cells, the SCP increased coverage of HLA-I and HLA-II immunopeptidomes by at least 2-fold compared to the Exploris + FAIMS.

### Reproducibility, variance, and dynamic range of HLA-I immunopeptidome analysis on the SCP

To accurately represent biological conditions and capture differences in low abundant clinically relevant antigens, low stochasticity, high sensitivity, wide dynamic range, and reproducible quantification is critical. Consistent with previous reports, we observed a ∼80% overlap of peptide IDs between technical replicates for samples run on the SCP and the Exploris + FAIMS across all input amounts (19, 26) (Figure 3A, Table S2 and 3). Figure 3B shows the overlap and uniqueness of HLA-I peptides observed on the instruments. At the 1e6 cell input level, only 5% of the 3057 HLA-I peptides identified are unique to the Exploris + FAIMS while 73% are unique to the SCP (Figure 3B, top panel). At 4e7 cell input level, 16% of the 19,477 HLA-I peptides observed are unique to the Exploris + FAIMS while 59% were only detected on the SCP (Figure 3B, bottom panel). Peptides observed in common in charge states +2 and +3 were found in the same charge states on the two instruments (Figure S6A). A small percentage (16%) of peptides were exclusively observed as +1 precursors on the SCP but +2 on the Exploris + FAIMS. Similarly, 12% of peptides were detected in both +1 and +2 charge states on the SCP and only in +2 on the Exploris + FAIMS (Figure S5A). Of note, around 25% of all HLA-I peptides identified using the SCP are observed in multiple charge states compared to only <10% on the Exploris + FAIMS (Figure S6B). The presence of multiple charge states per peptide provides additional confidence for peptide identification, which is particularly advantageous for immunopeptidome and other profiling applications such as post translational modifications that rely on single peptides.

**Figure 3:**
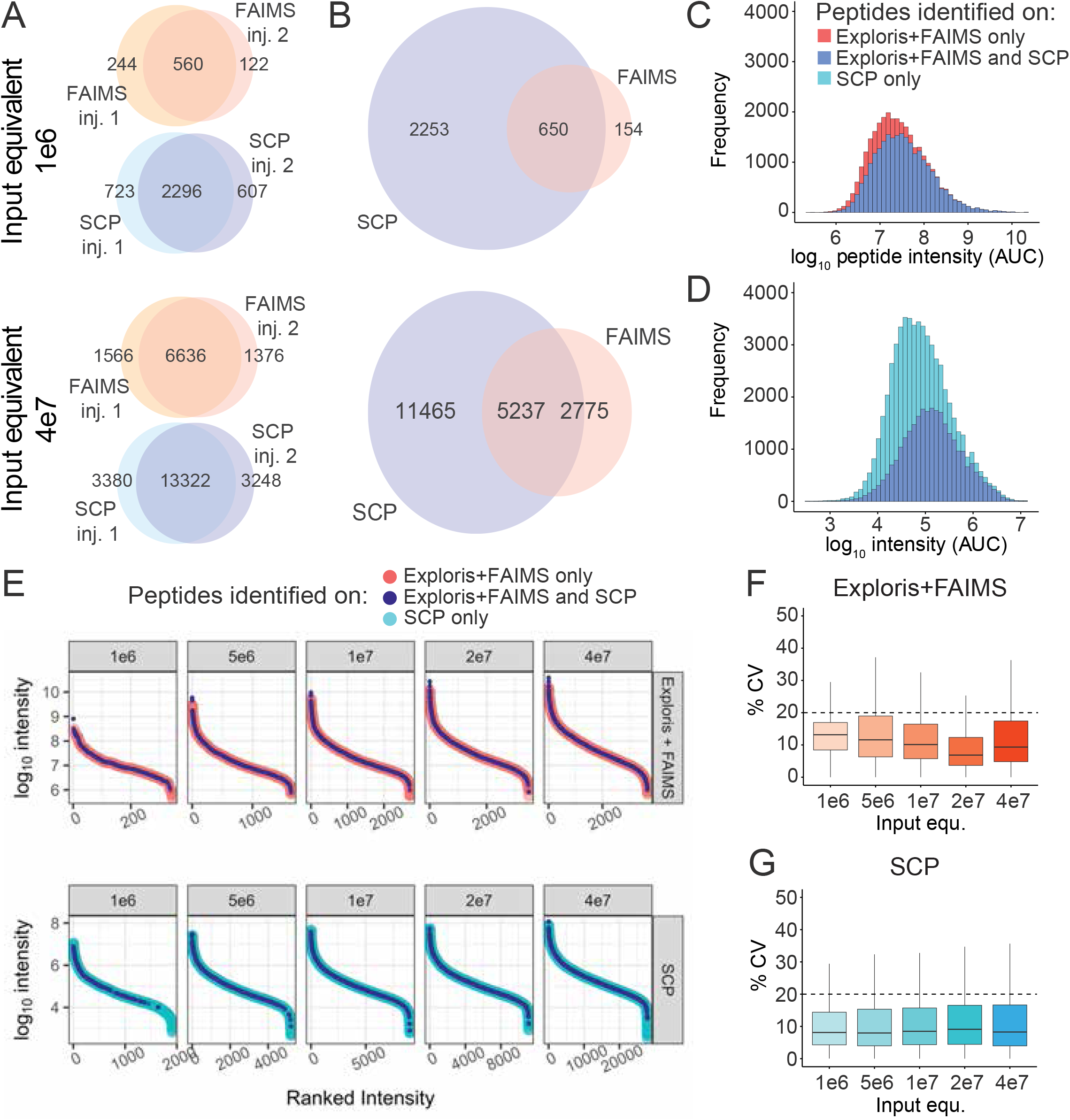
HLA-I peptide identifications on the timsTOF SCP show high reproducibility and extended dynamic range. A, Overlap of HLA-I peptides across multiple injections on the Exploris + FAIMS (FAIMS) or the timsTOF SCP at 1e6 or 4e7 A375 cell input equivalents. B, Overlap of HLA-I peptides between the Exploris + FAIMS (FAIMS) and the timsTOF SCP at 1e6 (upper panel) or 4e7 (lower panel) A375 cells. C, Log_10_ intensity (MS1 AUC) distribution of HLA-I peptides uniquely identified on the Exploris + FAIMS (pink) or both instruments (dark blue). D, Log_10_ intensity (MS1 AUC) distribution of HLA-I peptides uniquely identified on the timsTOF SCP (light blue) or both instruments (dark blue). E, Ranked peptide intensity at different cell input equivalents on the Exploris + FAIMS (upper panel) or the timsTOF SCP (lower panel). Respective intensity of peptides uniquely identified on the Exploris + FAIMS (pink), the timsTOF SCP (light blue) or both instruments (dark blue). F, Percent % CV of peptide intensity between technical replicates at different cell input equivalents on Exploris + FAIMS. Dashed line indicates 20% CV. G, % CV of peptide intensity between technical replicates identified on the timsTOF SCP at indicated input equivalents. Dashed line indicates 20% CV. AUC is area under the curve, %CV is coefficient of variation.

We hypothesized that >50% more unique HLA-peptides are identified on the SCP due to the instruments’ higher sensitivity and sampling speed (Figure 3B). Indeed, the median log_10_ precursor intensity of peptides uniquely identified on the SCP is lower compared to those observed in common (5.1 vs 4.9, Figure 3C-D). This is also illustrated by the extension of two orders of magnitude in overall dynamic range using the SCP compared to the Exploris + FAIMS (Figure 3E). The improved sampling of lower abundant peptides and the corresponding increase in dynamic range was observed at all input amounts on the SCP (Figure 3E). The median % coefficient of variation (%CV) on each instrument is <20% (Figure 3F-G), with a Pearson correlation of >0.8 between replicates across all input amounts (Figure S6A-B).

The results presented thus far were based on serially diluted A375 cells. As such, we directly enriched HLA-I peptides from low cell numbers to better understand the HLA-I immunopeptidome profiling depth of input limited samples. For this analysis we used the same input range as thes bulk experiments and acquired them with M10 on the SCP (Figure 4A and Table S4). Direct enrichment from 4e7 cells decreased HLA-I peptide identification by only 12% (i.e. 17,000 from bulk enrichment, 15,000 via direct IP). As expected, this decrease was amplified at lower cell input. We identified 700-900 HLA-I peptides from 1e6 cells, 2000-3000 from 5e6 cells, and 10,000 from 2e7 cells, respectively (Figure 4A, Table S4). Importantly, the immunopeptidome depth obtained by enrichment from as little as 1e6 A375 cells is sufficient to identify HLA-I peptides derived from cancer-testis antigen (CTA), and nuORFs (Figure 4B-C) (6, 34). Moreover, we observed CTA source protein derived peptides across all input levels, with ∼9 and ∼90 from 1e6 or 4e7 A375 cells, respectively (Figure 4B). Moreover, we detected ∼5 unique nuORF derived peptides from 1e6 A375 cells, with highest relative contribution from out-of-frame ORFs and 5’ uORFs across all sample input levels (Figure 4C).

**Figure 4:**
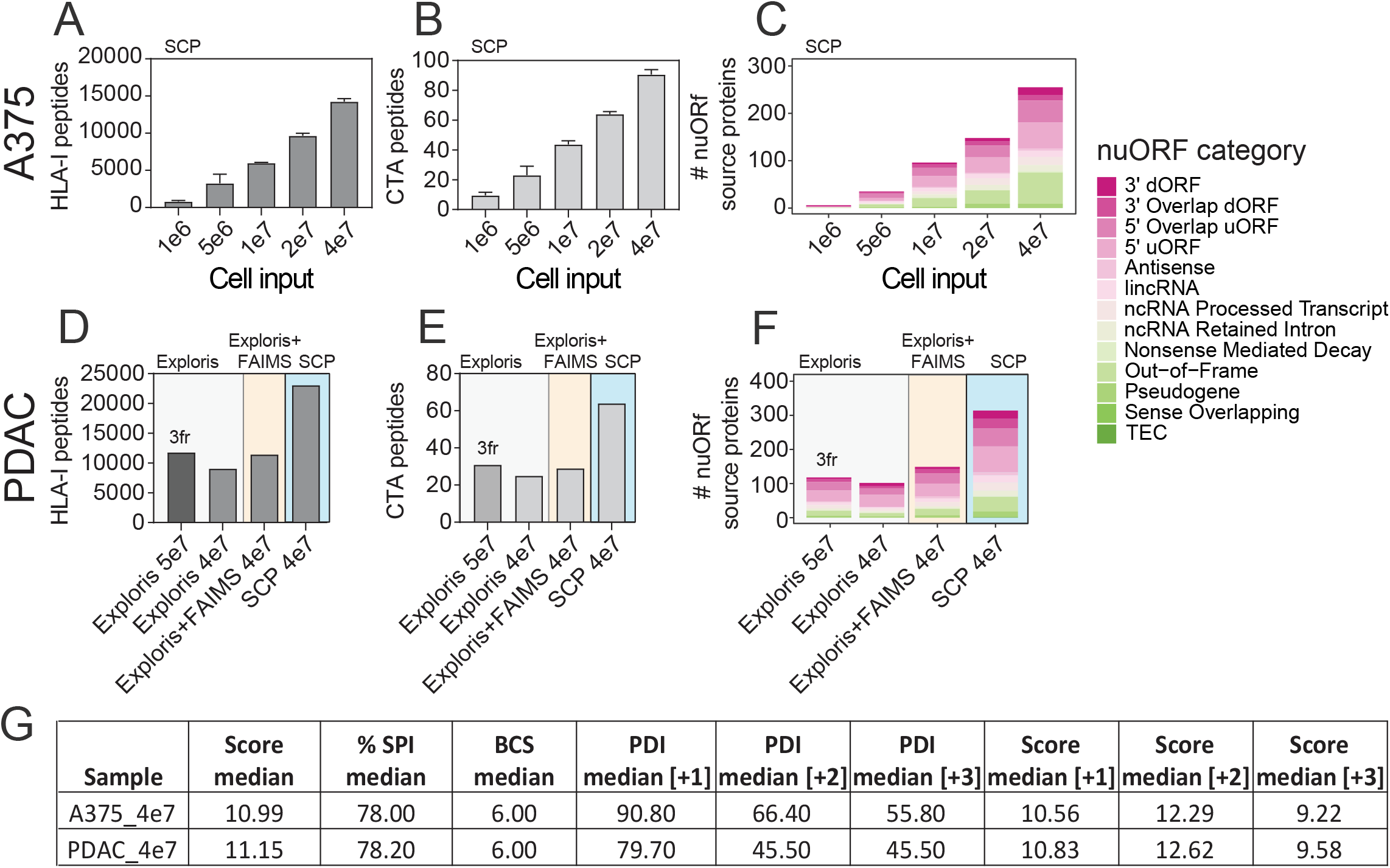
Single-shot acquisition of HLA-I peptides from low amounts of tumor derived cell lines enables detection of CTA and novel nuORFs derived peptides on the timsTOF SCP. A, Unique HLA-I peptides directly enriched from 1-40 million A375 cells as indicated by single-shot injections on timsTOF SCP. B, Peptides detected in panel A that map to CTA source proteins from CTdatabase. C, Unique nuORF source proteins contributing to HLA-I immunopeptidome of 1-40 million A375 cells. D, Unique HLA-I peptides identified from 40-50 million PDAC line in 3 offline stagetip fractions on the Exploris (Exploris, 3fr) or single-shot injections on Exploris ± FAIMS or the timsTOF SCP. E, Peptides detected in panel D that map to CTA source proteins from CTdatabase. F, Unique nuORF source proteins identified in a patient derived PDAC cell line with the acquisition schemes indicated in D. G, Quality metrics for HLA-I peptides from 4e7 A375 or PDAC samples including median score, % SPI, BCS and median PDI by charge state on the SCP. CTA is cancer testis antigens, nuORFs is unannotated open reading frames, PDAC is patient derived adenocarcinoma tumor cell, % SPI is percent scored peak intensity, BCS is backbone cleavage score, PDI is percent precursor dissociation intensity.

### Application of acquisition methods to samples with different HLA-I allele compositions

We next evaluated the M10 method that we optimized using the A375 cells on a sample with HLA alleles and peptide binding characteristics distinct from A375. For this we selected a patient derived PDAC line where we had sufficient input material to be able to evaluate both the Exploris + FAIMS and the SCP. We compared use of the single-shot M10 SCP method with single-shot injections ± FAIMS or brp fractionation (3 fr) without FAIMS on the Exploris (Table S8). Single-shot acquisition from 4e7 PDAC derived cells on the Exploris yielded 9080 HLA-I peptides without FAIMS and 11,460 with FAIMS. In contrast, 23,000 HLA-I peptides were identified on the SCP from 4e7 PDAC derived cells, an increase of > than 2 fold (Figure 4D). Use of off-line biochemical fractionation into 3 frs and single shot analysis using the Exploris only marginally increased the numbers of peptides identified versus single shot injections (11,810 vs. 11,460) while increasing manual sample handling and 3-fold increased data acquisition time, a trade-off that is impractical for large sample cohorts. Similarly, and consistent with observations in A375 cells, the SCP identifies 2-fold more CTA and nuORF derived HLA-I peptides (Figure 4E-F, Table S9).

While the median score, % SPI and BCS were comparable between the data collected on the Exploris and SCP for HLA-I peptides derived from the PDAC cells, we noted a >10% decrease in median PDI per charge state when compared to the A375 results (Figure 4G, Figure S7A). Specifically, PDI for +1, +2 and +3 charged precursors were reduced by 12%, 31% and 18%, respectively with +2 and +3 charge states affected the most (Figure 4G, Figure S8A-B). We concluded that the lower PDI on the SCP compared to Exploris + FAIMS (Figure 4G) was likely due to too low CE for optimal fragmentation of peptides with the HLA allele specific peptide-binding characteristics present in the PDAC cell line.

Aiming to identify an acquisition method on the SCP that could be universally applied to analyze patient cohorts with diverse HLA alleles, we revisited the CE parameters. Based on the observation that M10 particularly under-fragmented multiply charged peptides, we increased the CE at low IM 1/K0 = 0.6 Vs cm^-2^ from 10eV in method M10 to 15eV which we refer to as M11. This marginal increase in CE at low IM yielded comparable numbers of HLA-I peptides to the M10 method, but improved PDI for +2 and +3 charged precursors in melanoma samples (Figure S4B, S8A-D). We then tested both the M10 and M11 methods for analysis of the immunopeptidomes of primary melanoma tumors (n=5) from caucasian patients in another multi-allelic set of samples. The higher CE slope of M11 again increased PDI by 0.5-2.6% for +1, 13-25% for +2 and 17-35% for +3 precursors relative to M10 across all samples (Figure S9E). Importantly, the motifs of peptides identified with M10 and M11 revealed the same anchor residues (Figure S10A). Overall, we observed comparable numbers of peptides (8847 – 16,385) and scores (11.77 ± 5%) using M10 and M11 methods starting with just 15 mg cryopulverized tissue (Figures 5A, D, S8D-E). Moreover, we identified similar numbers of 20-60 CTA derived peptides and 100-300 unique nuORF peptides with both methods (Figure 5B-C, Table S11).

**Figure 5:**
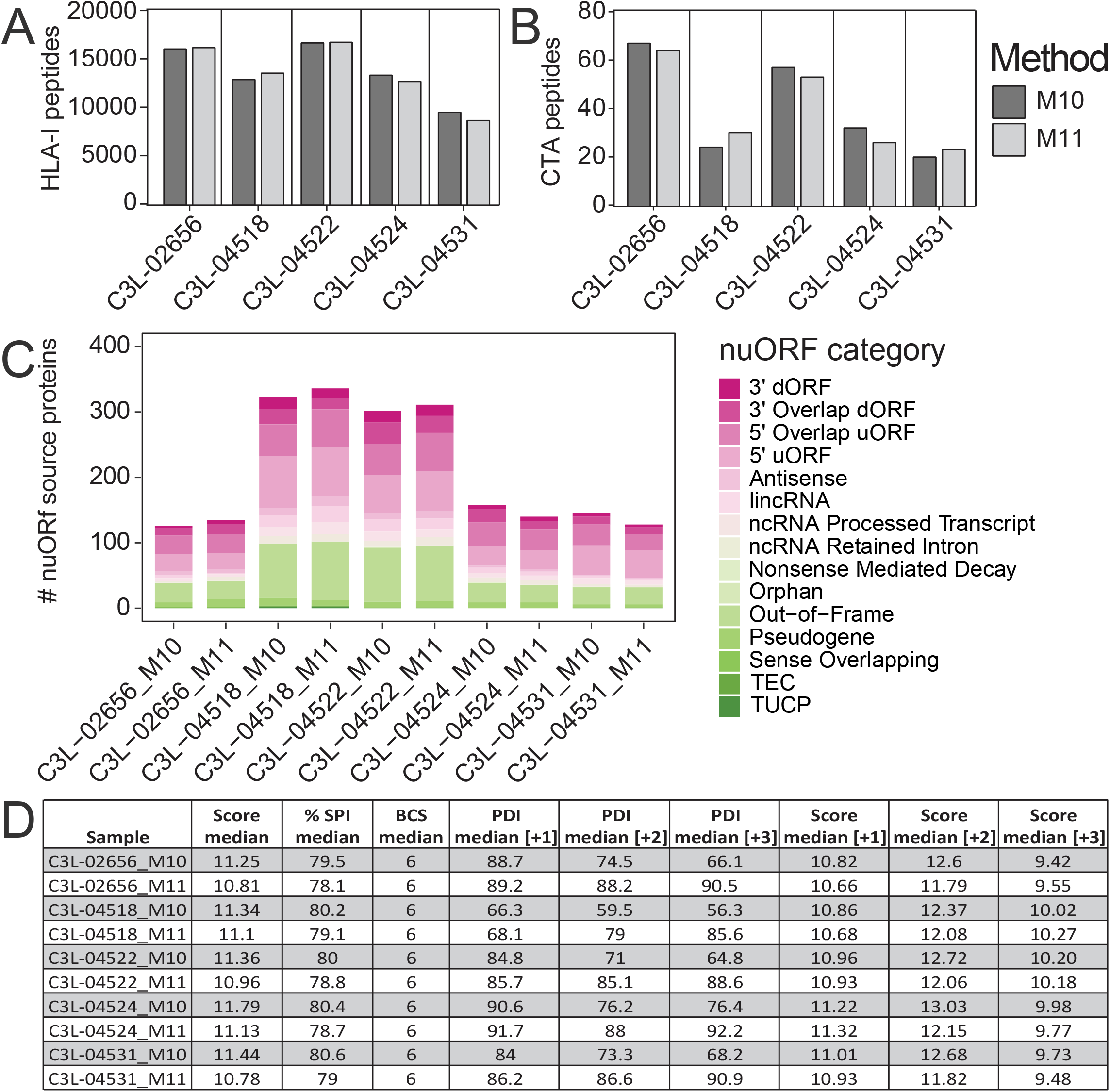
Single-shot DDA-PASEF of low-input primary melanoma tumors on the timsTOF SCP enables deep profiling of the HLA-I immunopeptidome and detection of CTA and nuORF derived peptides. A, Unique HLA-I peptides enriched from 15 mg wet weight primary melanoma tumors (estimated <1e7 cells) identified by single-shot injections on timsTOF SCP with M10-M11. B, Peptides detected in panel A that map CTA source proteins from CTdatabase. C, Unique nuORF source proteins represented in HLA-I immunopeptidome of the respective melanoma tumors. D, Quality metrics of HLA-I peptides from primary melanoma tumors including median score, % SPI, BCS, median PDI and scores by charge state using M10 or M11 on the SCP. CTA is cancer testis antigens, nuORFs is unannotated open reading frames, % SPI is percent scored peak intensity, BCS is backbone cleavage score, PDI is percent precursor dissociation intensity.

We next evaluated peptides exclusively identified by either M10 or M11 and observed an increase in singly charged precursors in M10. The slightly increased CE in M11 yielded more triply charged precursors with a slight shift in length towards 10-11aa as compared to 9aa in M10 (Figure S10B-C). We therefore speculate that higher CE is identifying longer (>9mer) and higher charged peptides because of improved fragmentation of these peptides.

In conclusion, we find that CE slopes presented here should be tested beforehand based on the anticipated sample or cohort specific HLA-I alleles and their peptide binding motifs, with M11 as our optimal method for immunopeptidome profiling from Caucasian populations.

## Discussion

One method that is frequently used to achieve deep immunopeptidome depth to enable detection of therapeutically relevant HLA antigens is offline orthogonal chromatographic separation. However, clinical sample availability is often limited, and offline fractionation does not offer the same benefits at lower input amounts and also significantly decreases sample throughput. To simultaneously improve immunopeptidome depth and increase analysis throughput for low-level samples, we evaluated and optimized acquisition parameters on the Bruker timsTOF SCP for HLA-I and HLA-II peptides. Through systematic evaluation of instrument parameters such as ion-mobility precursor selection polygon filter, collision energy settings, TIMS accumulation and ramp times and target intensity thresholds, we were able to optimize instrument parameters that greatly improved immunopeptidome depth. At higher input levels of 1e7 cells, we observe ∼10,000 HLA-I or 9,000 HLA-II peptides that included peptides derived from CTA and nuORF source proteins. Most importantly, method optimization enabled single shot immunopeptidome depth of up to 1000 HLA-I peptides from a direct IP of only 1e6 A375 cells and enhanced spectral quality as measured by score. The ability to characterize such low input samples also allows the direct detection of potential therapeutically relevant HLA peptides and facilitates immunopeptidome data collection from smaller and more difficult to obtain tumor biopsies.

Our systematic analysis found that multiple instrument method parameters contributed to the improved immunopeptidome depth. Specifically, we noted that extending the polygon filter placement to include singly charged precursors and decreasing the CE values had the largest impact on total peptide identifications (i.e. 35% increase compared to the default tryptic method, Figure 1D). The inclusion of singly charged precursors through adjustment of the polygon filter increased the number of charge states per HLA-I peptide by 15% thereby increasing the identification confidence through detection and fragmentation of the same peptide in multiple charge states. Surprisingly, a lower CE resulted in overall higher b and y ion pairs, which in turn increased the identification score even though the peptides appear to be somewhat under fragmented. While decreasing the CE did not impact the BCS, we observed different fragment ion type distributions produced on the Exploris vs. SCP. Specifically, we identified fewer internal fragment ions on the SCP compared to the Exploris + FAIMS and largely increased b and b/y fragment ion pairs, contributing to increased scores. We speculate that the high sensitivity of the SCP enables increased detection of the low abundant b/y fragment ion pairs formed as a result of decreased collision energy. Although the decreased CE slopes improve HLA-I peptide identifications and scores, we hypothesize that implementing precursor fragmentation in a charge state specific manner on the SCP, which is not currently possible with the version of instrument control software used in this study, will further improve peptide fragmentation to improve sequence coverage.

We also applied our optimized SCP analysis methods to profile the immunopeptidomes of a patient derived PDAC cell line and primary melanoma tumors identifying ∼22,000 HLA-I peptides from 4e7 PDAC cells and 8,847-16,3885 from 15 mg melanoma tissue. Interestingly, we noticed substantial differences in PDI between the initial method optimization (i.e. A375 cell line) and the subsequent application to patient-derived PDAC cell lines. Despite up to a 50% difference in PDI for multiply charged precursors, we observed comparable overall peptide scores, %SPI or the percent of peptide backbone cleavage in the PDAC cells vs. the A375 cells. Nonetheless, we evaluated a slightly higher CE slope in method M11 in an attempt to further improve fragmentation patterns. HLA-I peptide identifications and allele assignments were maintained relative to M10 in melanoma samples while improving PDI. The significant increase in precursor fragmentation for all charge states with M11 suggests that the use of the intermediate CE used in this method may be of more general use in the immunopeptidome analysis in larger cohorts that harbor large numbers of diverse HLA alleles. In addition, a minor fraction of peptides (less than 2%) unique to M11 were longer (>9mer) and present at higher charge state, therefore this method could benefit samples containing alleles that bind peptides containing basic amino acids. Therefore, for most comprehensive immunopeptidome depth and peptide representation, we recommend M11 for large scale experiments.

A limitation of this study is that only Caucasian patient samples were included in the melanoma tumor cohort, which might bias the observed allele distribution or peptide features for which we optimized instrument data acquisition parameters. Based on this we hypothesize that future studies with more comprehensive patient cohorts from diverse genetic backgrounds will provide further insights into improving immunopeptidome data acquisition parameters for diverse HLA allele populations. We believe that deep single shot immunopeptidome datasets generated on the SCP will improve HLA binding and presentation prediction algorithms that are presently trained using predominantly ion trap and Orbitrap data (26, 35–37). Similarly, CCS prediction has been shown to increase confidence in low abundant peptide identifications of complex samples, but so far have been mainly trained on tryptic peptides (20). We therefore speculate that the generation of comprehensive HLA-I and HLA-II datasets to train CCS prediction algorithms on the unique features of HLA peptides will greatly benefit analysis immunopeptidome data depth and help to improve the ability to directly identify low level clinically relevant HLA antigens, such as neoantigens. Finally, stochastic sampling by the MS systems and, possibly, background introduced by the antibody enrichment continues to limit overlap between repeated single-shot injections of HLA-I enriched samples to ca. 80%. We speculate that the improved dynamic range of the recently introduced high throughput TIMS cartridge (timsTOF HT) in combination with rapid data independent analysis PASEF acquisition could greatly benefit data reproducibility and more efficiently sample low abundant peptide species. In summary, we believe that the SCP method optimization and systematic analyses presented here will enable deep immunopeptidome profiling of large patient cohorts, which in turn will facilitate improvement to HLA peptide presentation and CCS prediction algorithms.

## Abbreviations

AGC: Automatic gain control
BCS: backbone cleavage score
brp: basic reversed phase
CCS: collisional cross-section
CE: collision energy
CTA: cancer-testis antigen
CV: compensation voltages
%CV: % coefficient of variation
Exploris + FAIMS: Thermo Fisher Orbitrap Exploris 480 + FAIMS ProTM
FDR: false discovery rate
HCD: Higher energy collisional dissociation
HLA-I: human leukocyte antigen class I
HLA-II: human leukocyte antigen class II
IM: Ion mobility
nuORFs: novel unannotated open reading frames
PASEF: Parallel accumulation-serial fragmentation
PDAC: pancreatic ductal adenocarcinoma tumor cell
%PDI: percent dissociated precursor intensity
PSM: Peptide spectrum match
SCP: Bruker timsTOF SCP
%SPI: percent scored peak intensity
SM: Spectrum Mill

## Acknowledgements

This work was supported in part by grants P01CA206978 to S.A.C from the NIH, U24CA270823, U01CA271402 to S.A.C, and U24-CA271075 to D.R.M. from National Cancer Institute (NCI) Clinical Proteomic Tumor Analysis Consortium program, as well as a grant from the Swiss National Science Foundation (SNF) grant CRSII5_186405 to N.D.U. and S.A.C., and from the Dr. Miriam and Sheldon G. Adelson Medical Research Foundation to N.D.U. and S.A.C.. N.D.U. is a recipient of a SPARC Award to from the Broad Institute of MIT & Harvard (#800373) that partially supported this work.

We thank Hasmik Keshishian and Matt Willetts for assistance with mass spectrometry instrumentation and methods, Khoi Munchic Pham and D R Mani for assistance with Fragpipe analysis on Terra, Jackson White and Joseph Allen for assistance with melanoma tumor sample processing, Andrew Aguirre, William A Freed-Pastor, Zack Ely and Tyler Jacks for access to the patient organoid derived pancreatic cancer cell line, and Michael Gillette for melanoma tumor sample procurement.

## Conflict of Interest

S.A.C. is a member of the scientific advisory boards of Kymera, PTM BioLabs, Seer and PrognomIQ.

A.S.V.J. is an employee of Bruker.

## Data availability

The raw mass spectrometry data have been deposited in the public proteomics repository MassIVE for reviewer access. This data will be made public upon acceptance of the manuscript.

## Notes

### Competing Interest Statement

The authors have declared no competing interest.

### Summary of Updates

Data availability section updated as follows: The raw mass spectrometry data have been deposited in the public proteomics repository MassIVE for reviewer access. This data will be made public upon acceptance of the manuscript.

