## Supplementary Figures for "Sensitive, high-throughput HLA-I and HLA-II immunopeptidomics using parallel accumulation-serial fragmentation mass spectrometry"

A

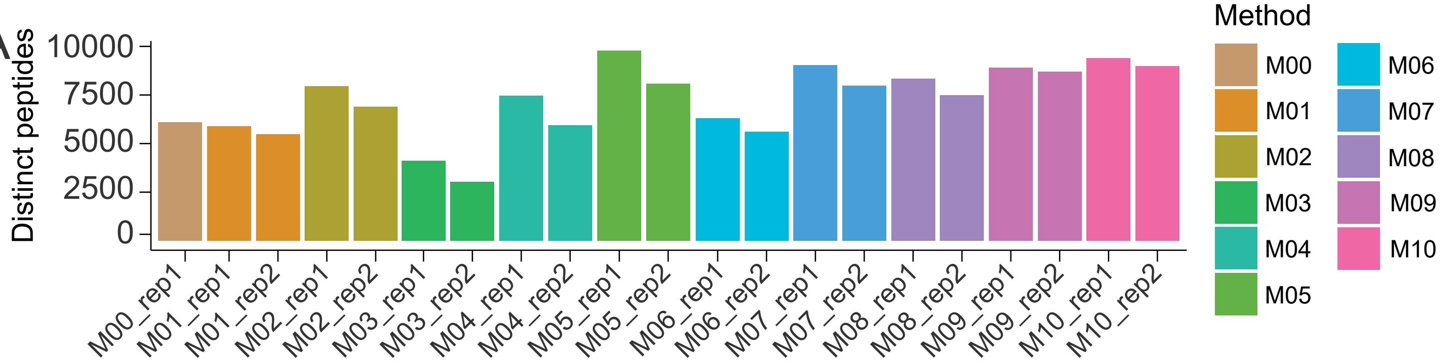


B

| directory | CE | HLA-I peptides | Score median | % SPI median | % BCS median | PDI median [+1] | PDI median [+2] | PDI median [+3] | Score median [+1] | Score median [+2] | Score median [+3] |
|---|---|---|---|---|---|---|---|---|---|---|---|
| M00 - rep1 | 20-59 | 6447 | 8.2 | 68.7 | 6.0 | 67.3 | 93.2 | 95.7 | 12.1 | 8.2 | 7.0 |
| M01 - rep1 | 20-59 | 6188 | 8.6 | 70.3 | 6.0 | 69.9 | 93.4 | 95.6 | 9.4 | 8.7 | 7.3 |
| M01 - rep2 | 20-59 | 5726 | 8.5 | 70.0 | 6.0 | 76.2 | 93.9 | 95.9 | 8.3 | 8.7 | 7.2 |
| M02 - rep1 | 20-59 | 8361 | 8.7 | 70.8 | 6.0 | 95.3 | 93.2 | 95.6 | 8.8 | 8.7 | 7.3 |
| M02 - rep2 | 20-59 | 7199 | 8.6 | 70.8 | 6.0 | 96.4 | 93.8 | 95.7 | 8.7 | 8.7 | 7.1 |
| M03 - rep1 | 30-65 | 4438 | 7.3 | 68.3 | 6.0 | 98.1 | 97.0 | 97.7 | 8.1 | 6.9 | 5.8 |
| M03 - rep2 | 30-65 | 3343 | 7.1 | 67.9 | 6.0 | 98.5 | 97.1 | 97.7 | 7.6 | 6.8 | 5.5 |
| M04 - rep1 | Stepped | 7791 | 8.4 | 70.0 | 6.0 | 98.0 | 93.2 | 95.7 | 8.0 | 8.6 | 7.2 |
| M04 - rep2 | Stepped | 6244 | 8.3 | 70.1 | 6.0 | 98.4 | 93.9 | 95.7 | 8.1 | 8.5 | 7.1 |
| M05 - rep1 | 10-55 | 10130 | 9.7 | 72.7 | 6.0 | 91.1 | 69.7 | 65.1 | 9.2 | 10.3 | 8.0 |
| M05 - rep2 | 10-55 | 8409 | 9.9 | 73.4 | 6.0 | 92.6 | 70.3 | 66.0 | 9.3 | 10.3 | 8.0 |
| M06 - rep1 | 20-59 | 6538 | 8.2 | 69.0 | 6.0 | 96.0 | 92.9 | 95.4 | 8.5 | 8.1 | 6.8 |
| M06 - rep2 | 20-59 | 6024 | 8.2 | 69.1 | 6.0 | 95.9 | 93.5 | 95.5 | 8.5 | 8.1 | 6.7 |
| M07 - rep1 | 20-59 | 9468 | 8.8 | 71.2 | 6.0 | 95.5 | 92.9 | 95.2 | 8.9 | 8.7 | 7.5 |
| M07 - rep2 | 20-59 | 8292 | 8.8 | 71.2 | 6.0 | 95.7 | 93.4 | 95.7 | 9.1 | 8.7 | 7.3 |
| M08 - rep1 | 20-59 | 8519 | 9.9 | 75.7 | 6.0 | 96.0 | 93.8 | 95.6 | 9.8 | 10.0 | 8.1 |
| M08 - rep2 | 20-59 | 7576 | 9.9 | 76.0 | 6.0 | 96.1 | 94.1 | 95.9 | 9.7 | 10.0 | 8.2 |
| M09 - rep1 | 10-55 | 9162 | 10.1 | 73.9 | 6.0 | 91.9 | 71.0 | 68.0 | 9.4 | 10.5 | 8.0 |
| M09 - rep2 | 10-55 | 9063 | 10.0 | 73.9 | 6.0 | 91.5 | 70.0 | 65.1 | 9.4 | 10.5 | 8.0 |
| M10 - rep1 | 10-55 | 9493 | 11.3 | 78.4 | 6.0 | 92.1 | 70.7 | 64.2 | 10.5 | 12.0 | 8.8 |
| M10 - rep2 | 10-55 | 9001 | 11.4 | 78.5 | 6.0 | 91.5 | 69.2 | 62.6 | 10.3 | 12.1 | 9.0 |

C D E

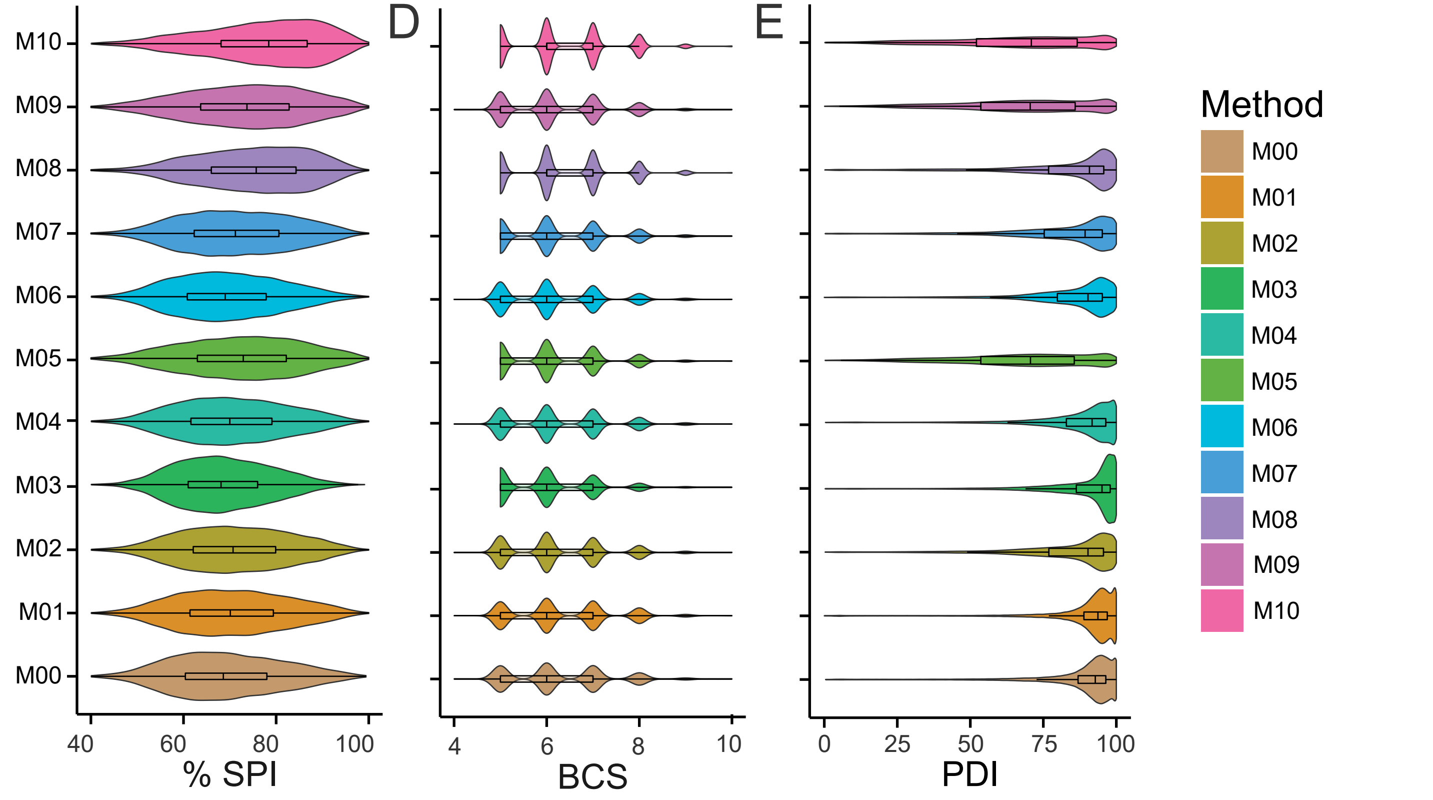

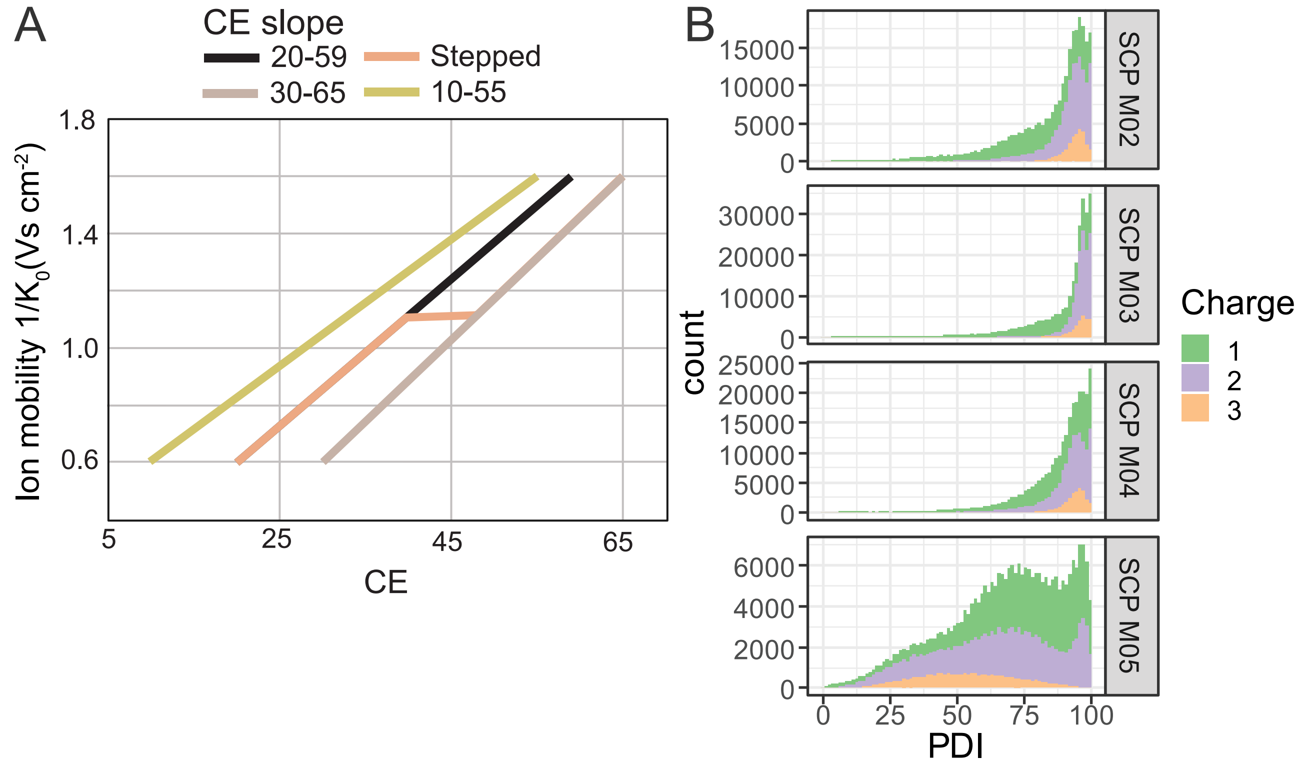
A
CE slope
20-59
Stepped
30-65
10-55
Ion mobility 1/$K_0$(Vs $cm^{-2}$)
1.8
1.4
1.0
0.6
5
25
45
65
CE
B
count
15000
10000
5000
0
30000
20000
10000
25000
20000
15000
10000
6000
4000
2000
SCP M02
SCP M03
SCP M04
SCP M05
0
25
50
75
100
PDI
Charge
1
2
3

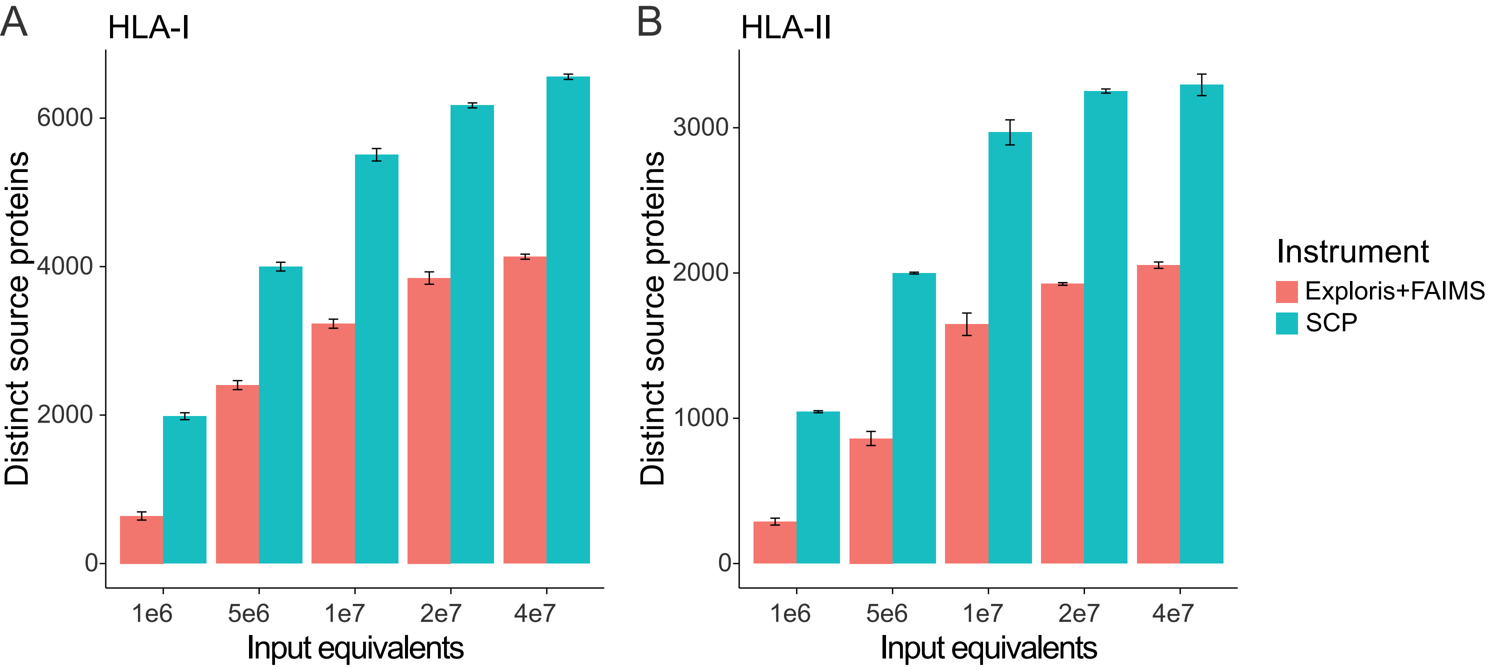

A
HLA-I
B
HLA-II
Distinct source proteins
0
2000
4000
6000
0
1000
2000
3000
1e6
5e6
1e7
2e7
4e7
Input equivalents
Instrument
Exploris+FAIMS
SCP

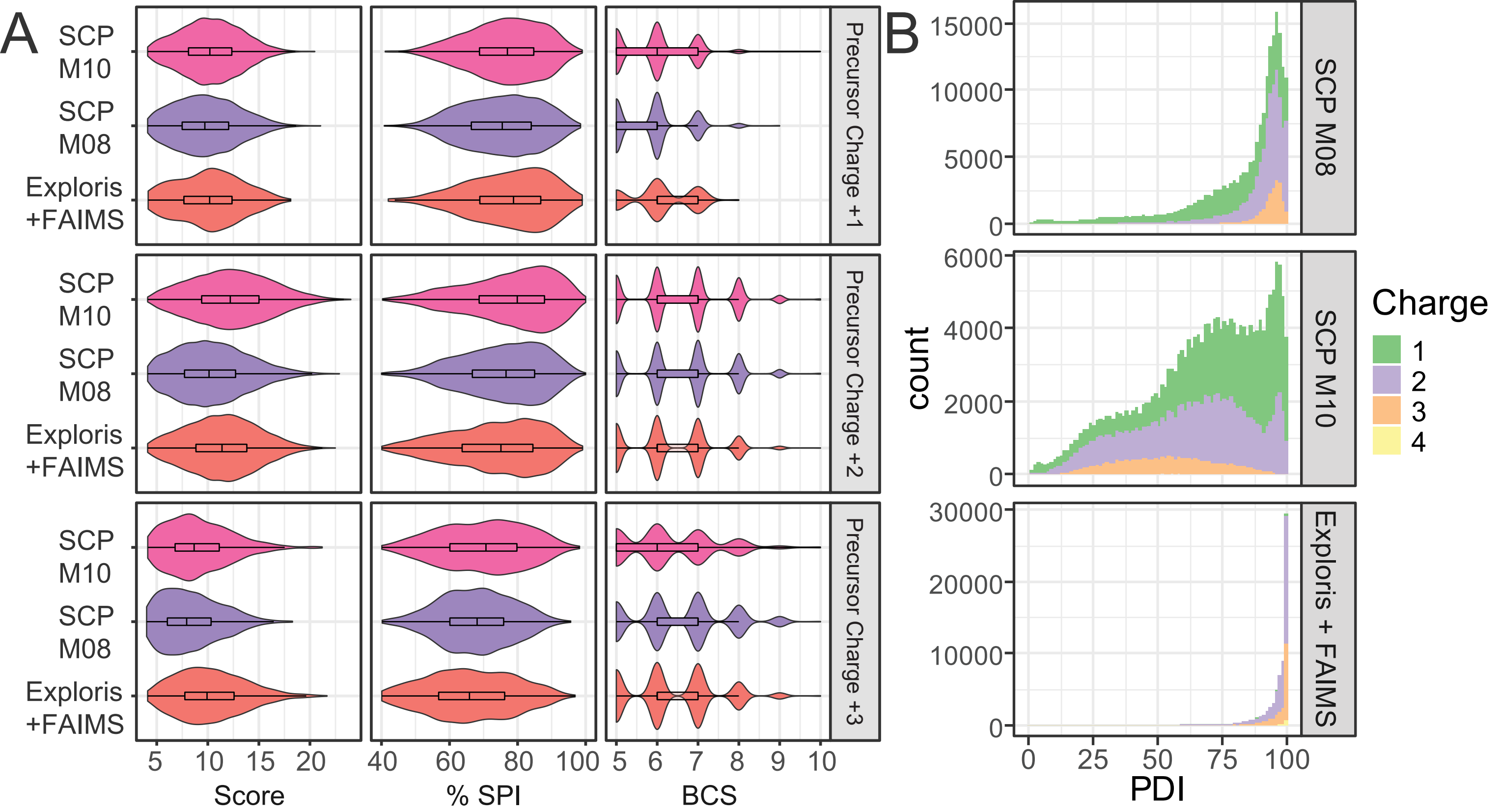


C

| Sample | Score median | % SPI median | % BCS median | PDI median [+1] | PDI median [+2] | PDI median [+3] | PDI median [+4] |
|---|---|---|---|---|---|---|---|
| SCP M10 | 11.3 | 78.4 | 6.0 | 92.1 | 70.7 | 64.2 | |
| SCP M08 | 9.9 | 75.7 | 6.0 | 96.0 | 93.8 | 95.6 | |
| Exploris+FAIMS | 10.63 | 75.9 | 6.0 | | 98 | 98.1 | 97.6 |

A

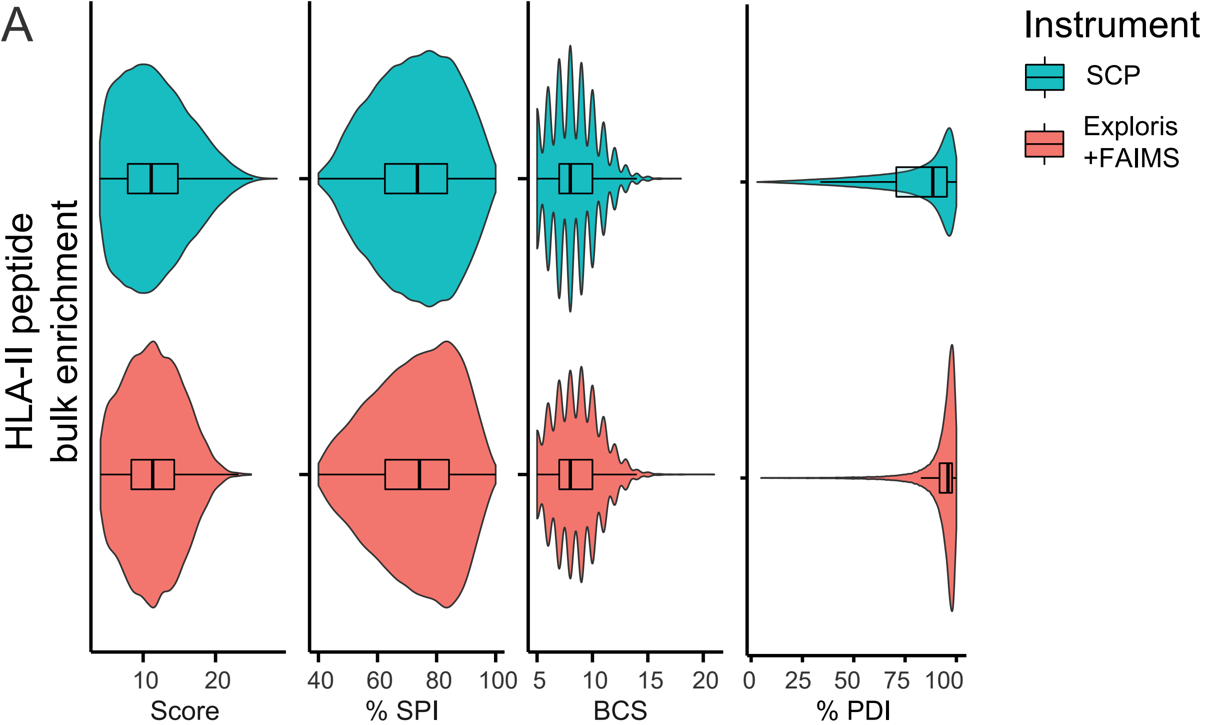


B

| Sample | Score median | % SPI median | % BCS median | PDI median [+2] | PDI median [+3] | PDI median [+4] | PDI median [+5] |
|---|---|---|---|---|---|---|---|
| SCP | 10.61 | 71.1 | 8 | 75.2 | 91.8 | 95.7 | 96.75 |
| Exploris+FAIMS | 10.85 | 74.9 | 8.5 | 98.8 | 98.9 | 98.9 | 98.6 |

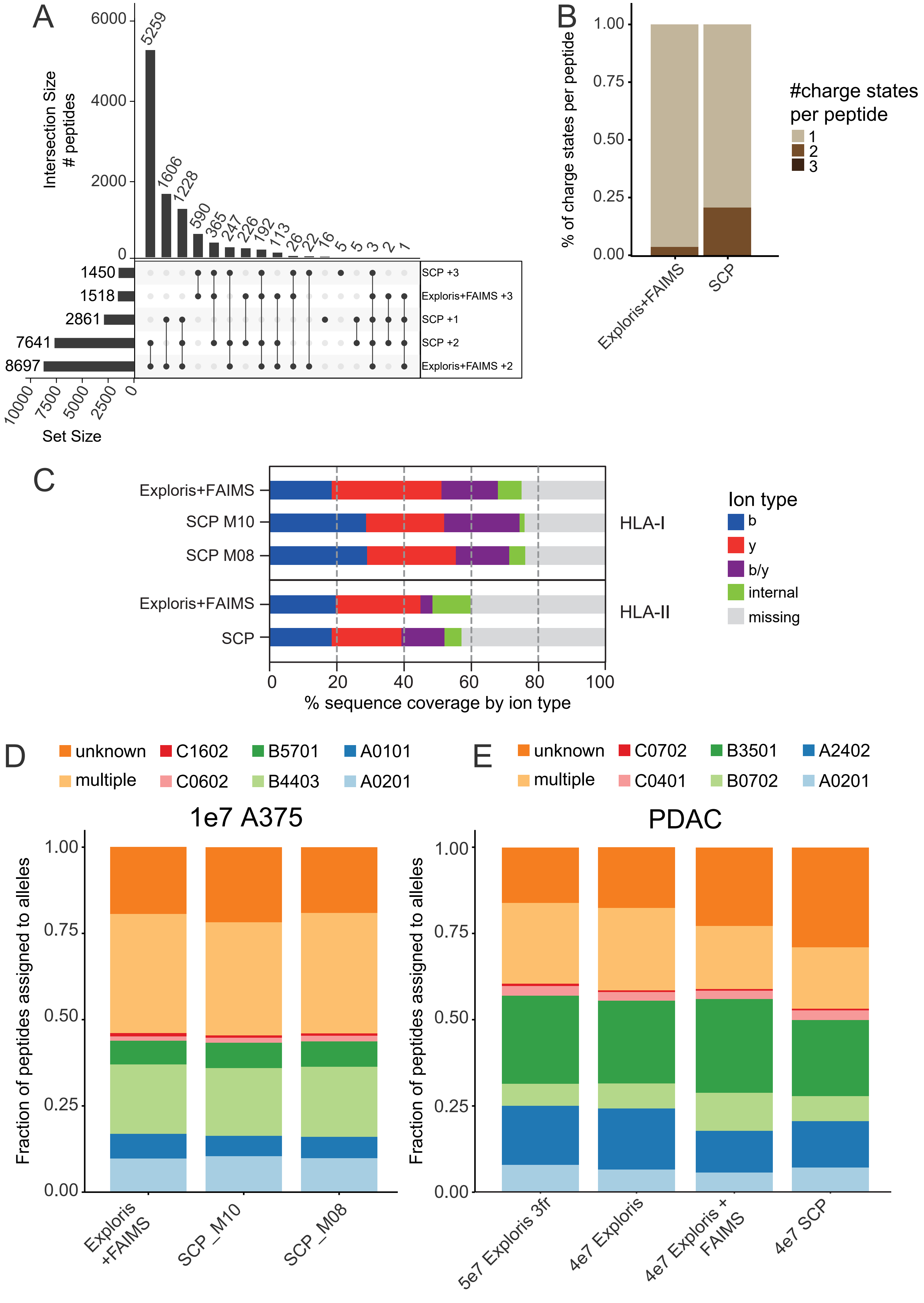
A
Intersection Size
### peptides
6000
4000
2000
0
5259
1606
1228
590
365
247
226
192
113
26
22
16
5
5
3
2
1
1450
1518
2861
7641
8697
SCP +3
Exploris+FAIMS +3
SCP +1
SCP +2
Exploris+FAIMS +2
10000
7500
5000
2500
0
Set Size
B
% of charge states per peptide
1.00
0.75
0.50
0.25
0.00
Exploris+FAIMS
SCP
#charge states per peptide
1
2
3
C
Exploris+FAIMS
SCP M10
SCP M08
Exploris+FAIMS
SCP
HLA-I
HLA-II
0
20
40
60
80
100
% sequence coverage by ion type
Ion type
b
y
b/y
internal
missing
D
unknown
C1602
B5701
A0101
multiple
C0602
B4403
A0201
1e7 A375
Fraction of peptides assigned to alleles
1.00
0.75
0.50
0.25
0.00
Exploris +FAIMS
SCP_M10
SCP_M08
E
unknown
C0702
B3501
A2402
multiple
C0401
B0702
A0201
PDAC
Fraction of peptides assigned to alleles
1.00
0.75
0.50
0.25
0.00
5e7 Exploris 3fr
4e7 Exploris
4e7 Exploris + FAIMS
4e7 SCP

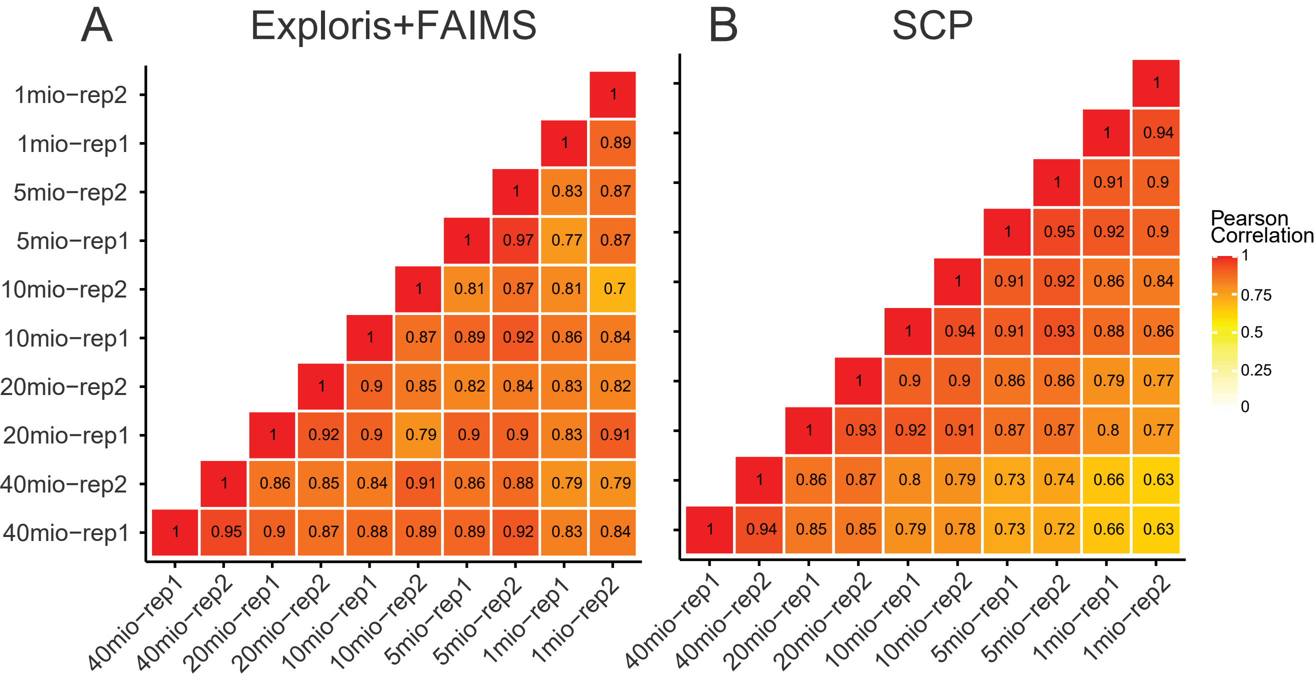
A
Exploris+FAIMS
B
SCP
1mio−rep2
1mio−rep1
5mio−rep2
5mio−rep1
10mio−rep2
10mio−rep1
20mio−rep2
20mio−rep1
40mio−rep2
40mio−rep1
40mio−rep1
40mio−rep2
20mio−rep1
20mio−rep2
10mio−rep1
10mio−rep2
5mio−rep1
5mio−rep2
1mio−rep1
1mio−rep2
1
1 0.89
1 0.83 0.87
1 0.97 0.77 0.87
1 0.81 0.87 0.81 0.7
1 0.87 0.89 0.92 0.86 0.84
1 0.9 0.85 0.82 0.84 0.83 0.82
1 0.92 0.9 0.79 0.9 0.9 0.83 0.91
1 0.86 0.85 0.84 0.91 0.86 0.88 0.79 0.79
1 0.95 0.9 0.87 0.88 0.89 0.89 0.92 0.83 0.84
1
1 0.94
1 0.91 0.9
1 0.95 0.92 0.9
1 0.91 0.92 0.86 0.84
1 0.94 0.91 0.93 0.88 0.86
1 0.9 0.9 0.86 0.86 0.79 0.77
1 0.93 0.92 0.91 0.87 0.87 0.8 0.77
1 0.86 0.87 0.8 0.79 0.73 0.74 0.66 0.63
1 0.94 0.85 0.85 0.79 0.78 0.73 0.72 0.66 0.63
Pearson Correlation
1
0.75
0.5
0.25
0

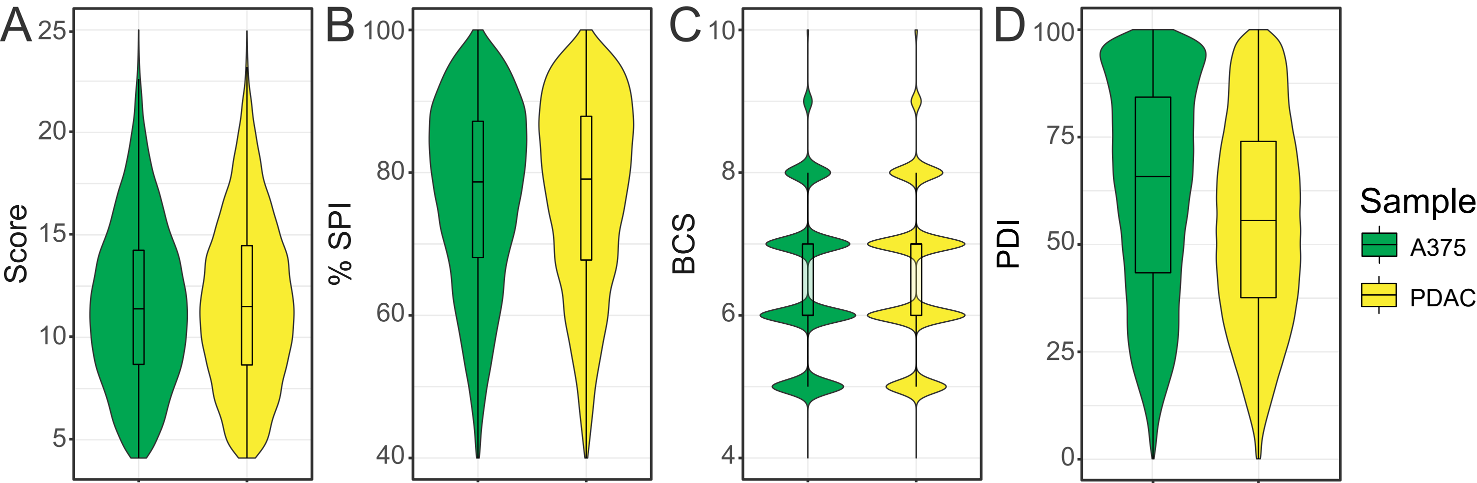
A
Score
25
20
15
10
5
B
% SPI
100
80
60
40
C
BCS
10
8
6
4
D
PDI
100
75
50
25
0
Sample
A375
PDAC

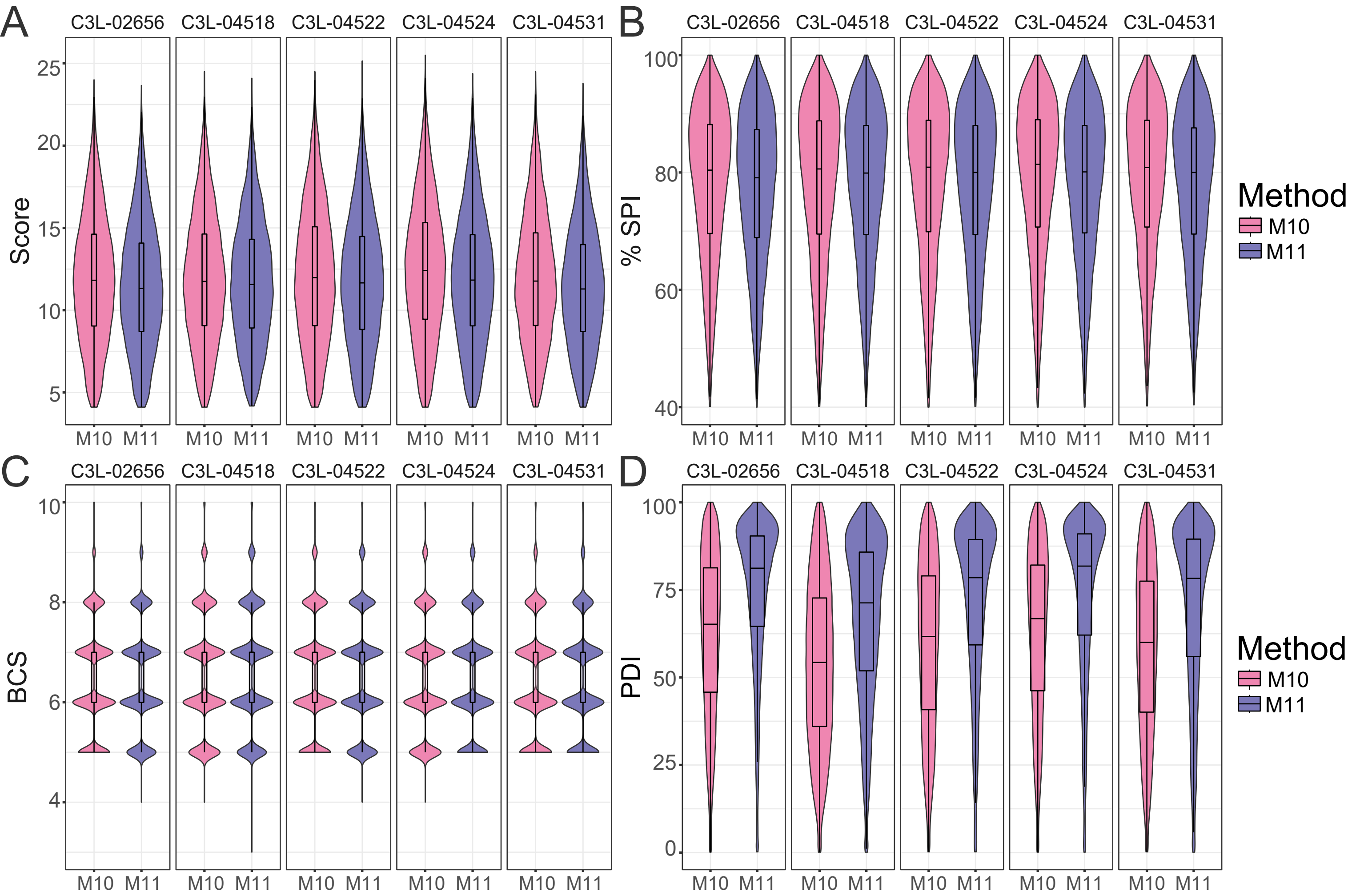

A
B
C
D
C3L-02656
C3L-04518
C3L-04522
C3L-04524
C3L-04531
Score
% SPI
BCS
PDI
M10
M11
Method

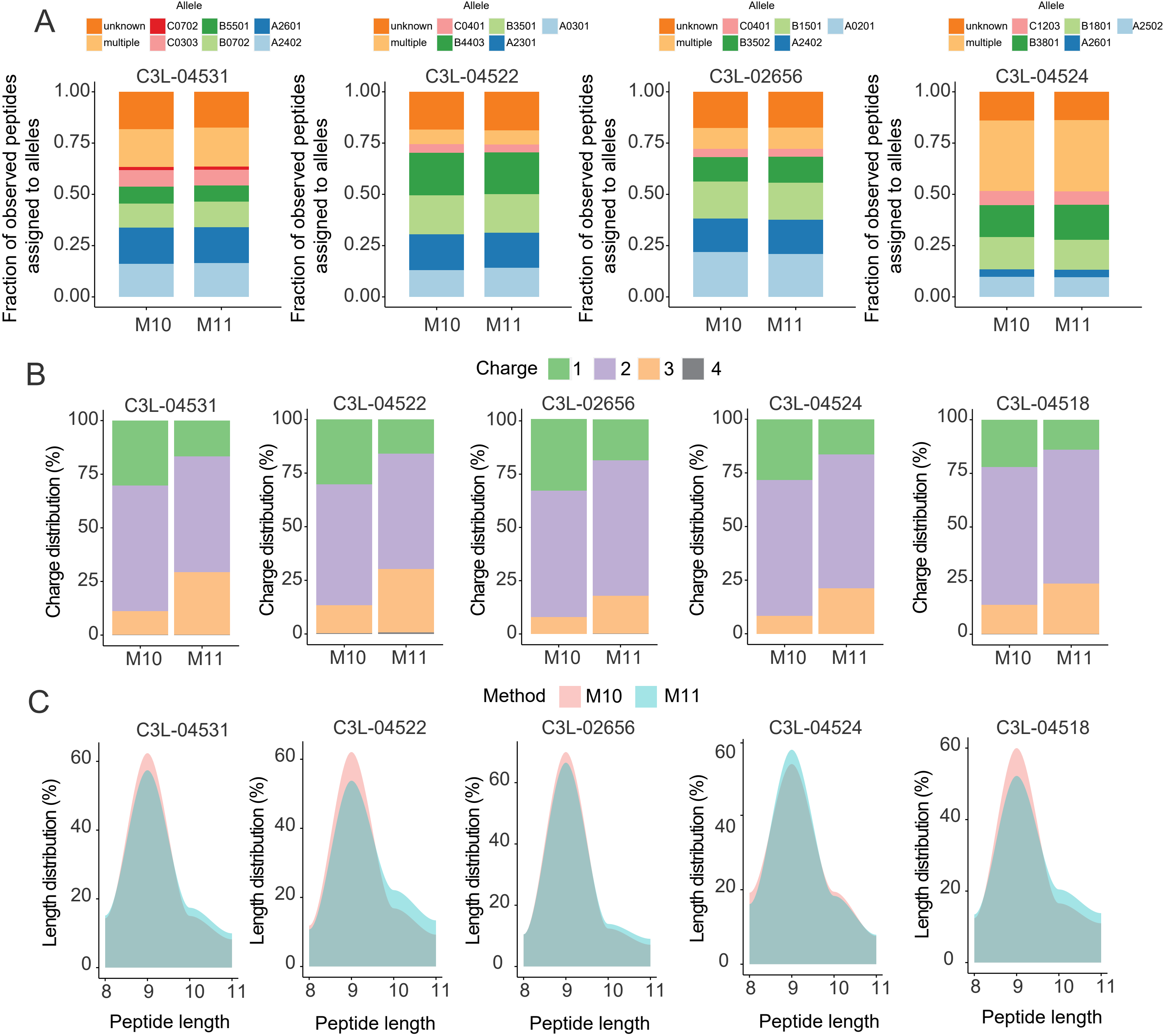
A
Allele
unknown
multiple
C0702
C0303
B5501
B0702
A2601
A2402
C3L-04531
Fraction of observed peptides assigned to alleles
1.00
0.75
0.50
0.25
0.00
M10
M11
Allele
unknown
multiple
C0401
B4403
B3501
A2301
A0301
C3L-04522
Allele
unknown
multiple
C0401
B3502
B1501
A2402
A0201
C3L-02656
Allele
unknown
multiple
C1203
B3801
B1801
A2601
A2502
C3L-04524
B
Charge
1
2
3
4
C3L-04531
C3L-04522
C3L-02656
C3L-04524
C3L-04518
Charge distribution (%)
100
75
50
25
0
M10
M11
C
Method
M10
M11
C3L-04531
C3L-04522
C3L-02656
C3L-04524
C3L-04518
Length distribution (%)
60
40
20
0
Peptide length
8
9
10
11
