## Supplementary Figure legends for "Sensitive, high-throughput HLA-I and HLA-II immunopeptidomics using parallel accumulation-serial fragmentation mass spectrometry"

### Supplementary Table Legends

**Supplementary Table Meta:** Metadata to Supplementary tables including patient specific HLA typing.

**Supplementary Table 1**: HLA-I peptide identifications from HLA-I method testing on the timsTOF SCP using A375 bulk digests at an input equivalent of 1e7 cells with acquisition methods M00-M10.

**Supplementary Table 2:** HLA-I peptide identifications from bulk titration on Exploris + FAIMS using A375 bulk digests.

**Supplementary Table 3:** HLA-I peptide identifications from bulk titration on timsTOF SCP with M10 using A375 bulk digests.

**Supplementary Table 4:** HLA-I peptide identifications from low input A375 cell HLA-I enrichment on timsTOF SCP with M10.

**Supplementary Table 5:** HLA-II peptide identifications from bulk titration on Exploris + FAIMS using A375 bulk digests.

**Supplementary Table 6:** HLA-II peptide identifications from bulk titration on timsTOF SCP using A375 bulk digests.

**Supplementary Table 7:** HLA-I peptide identifications from pancreatic ductal adenocarcinoma tumor cell (PDAC) line enrichments on timsTOF SCP with M10.

**Supplementary Table 8:** Novel unannotated open reading frames (nuORFs) identified from low-input HLA-I A375 and PDAC enrichments on the timsTOF SCP with M10.

**Supplementary Table 9:** HLA-I peptide identifications from low-input primary melanoma tumor on the timsTOF SCP with acquisition methods M10 and M11.

**Supplementary Table 10:** nuORFs identified from low-input primary melanoma tumor on the timsTOF SCP with acquisition methods M10 and M11.

### Supplemental Figure legends

**Supplemental Figure S1: Quality metrics of HLA-I peptides from A375 cells identified with different acquisition methods on the timsTOF SCP.**

A, Unique A375 HLA-I peptides identified from per injection replicate for M00-M10 on the timsTOF SCP.

B, Quality metrics for HLA-I peptides from A375 cells including median score, % SPI, BCS, PDI and scores by charge state using M00-M10.

C, Distribution of % SPI of A375 HLA-I peptides identified with M00-M10.

D, BCS distribution of HLA-I peptides identified with M00-M10.

E, PDI across all identified HLA-I peptides using M00-M10. % SPI is percent scored peak intensity, BCS is backbone cleavage score, PDI is percent precursor dissociation intensity.

**Supplemental Figure 2: CE dependency to IM impacts HLA-I peptide fragmentation on the SCP.**

A, Representative CE slopes 20-59 (M00-M02, M06-M08), 30-65 (M03), 10-55 (M05, M09-M10) from 1/K0 = 0.6 Vs $cm^{-2}$ to 1/K0 = 1.6 Vs $cm^{-2}$ or stepped (20-40 from 1/K0 = 0.6-1.1 Vs $cm^{-2}$ and 48-65 from 1/K0 = 1.1-1.6 Vs $cm^{-2}$).

B, PDI on the timsTOF SCP with M02-M05 by charge state. CE is collisional energy, PDI is percent precursor dissociation intensity.

**Supplemental Figure 3: Single-shot acquisition of HLA-I and HLA-II peptides on the timsTOF SCP increases source protein coverage >1.5-fold compared to Exploris + FAIMS.**

A, Unique source proteins represented in HLA-I immunopeptidomes by single injections on Exploris + FAIMS (red) and timsTOF SCP (blue). Mean and standard deviation is shown.

B, Unique source proteins represented in HLA-II immunopeptidomes by single injections on Exploris + FAIMS (red) and timsTOF SCP (blue). Mean and standard deviation is shown.

**Supplemental Figure 4: Quality metrics of HLA-I peptides identified on the timsTOF SCP upon CE alterations and the Exploris + FAIMS.**

A, Distributions of score, % SPI and BCS of HLA-I peptides on the timsTOF SCP with M10 (pink), M08 (purple) or the Exploris + FAIMS (orange) by charge state.

B, PDI on the timsTOF SCP with M08, M10 or the Exploris + FAIMS by charge state.

C, Quality metrics for HLA-I peptides including median score, % SPI, BCS, median PDI by charge state for the timsTOF SCP (M10 and M08) or the Exploris + FAIMS. CE is collisional energy, % SPI is percent scored peak intensity, BCS is backbone cleavage score, PDI is percent precursor dissociation intensity.

**Supplemental Figure 5: Quality metrics of HLA-II peptides identified on the timsTOF SCP and the Exploris + FAIMS are highly comparable.**

A, Distributions of score, % SPI, BCS and PDI of HLA-II peptides identified on the timsTOF SCP and Exploris + FAIMS.

B, Quality metrics for HLA-II peptides including median score, % SPI, BCS, median PDI by charge state for the SCP or the Exploris + FAIMS. % SPI is percent scored peak intensity, BCS is backbone cleavage score, PDI is percent precursor dissociation intensity.

**Supplemental Figure 6: Charge state distribution, sequence coverage by ion type and allele assignments of HLA-I and HLA-II peptides identified on the timsTOF SCP or Exploris ± FAIMS.**

A, Overlap of HLA-I peptides identified on both instruments in charge states 1, 2 or 3 on the timsTOF SCP or in charge states 2 and 3 on the Exploris + FAIMS.

B, Percentage of commonly identified HLA-I peptides on both the timsTOF SCP or Exploris + FAIMS in a single (beige), two (brown) or three (dark brown) charge states.

C, Percent sequence coverage by ion type for HLA-I and HLA-II peptides from the timsTOF SCP or the Exploris + FAIMS. b-ions in blue, y-ions in red, b/y-ion pairs in purple, internal-ions in green and missing/unassigned-ions in gray.

D, Allele assignment of 8-11 aa long HLA-I peptides identified on Exploris + FAIMS or SCP (M10 and M08), filtered for HLAthena rank <0.5 in 1e7 A375 cells.

E, Allele assignment of 8-11 aa long HLA-I peptides identified from the PDAC cell line across 3 fractions (3fr) or single-shot injections on the Exploris, Exploris + FAIMS and the timsTOF SCP. Data is filtered for HLAthena rank <0.5. PDAC is patient derived adenocarcinoma cells.

**Supplemental Figure 7: HLA-I analysis on the timsTOF SCP shows comparable reproducibility to Exploris + FAIMS.**

A, Intensity correlation of HLA-I peptides between technical replicates on the Exploris + FAIMS.

B, Intensity correlation between technical replicates of HLA-I peptides identified on the timsTOF SCP.

**Supplemental Figure 8: PDI of HLA-I peptides identified on the timsTOF SCP differs between A375 and PDAC cell lines.**

Distributions of score (A), %SPI (B), BCS (C), PDI (D) of HLA-I peptides from A375 cells (green) or PDAC line (yellow) on the timsTOF SCP. PDAC is patient derived adenocarcinoma cells.

**Supplemental Figure 9: CE alterations on the SCP improves PDI without impacting scores of HLA-I peptides from primary melanoma tumors.**

Score (A), % SPI (B), BCS (C), PDI (D) distributions of HLA-I peptides from primary melanoma tumors upon CE alterations (M10 or M11) acquired on the SCP.

E, Quality metrics for HLA-I peptides from primary melanoma tumors including median score, % SPI, BCS, PDI or score by charge state for the timsTOF SCP. CE is collisional energy, PDI is percent precursor dissociation intensity, % SPI is percent scored peak intensity, BCS is backbone cleavage score.

**Supplemental Figure 10: CE alterations on the timsTOF SCP impact charge state distribution of identified HLA-I peptides from primary melanoma tumors.**

A, Allele assignment of HLA-I 8-11 mers identified on the timsTOF SCP with either M10 or M11, filtered for HLAthena rank <0.5 in primary melanoma tumors.

B, Charge state distribution of HLA-I peptides from M10 or M11 on the timsTOF SCP.

C, Peptide length distribution across identified HLA-I peptides on the timsTOF SCP with M10 or M11 in primary melanoma tumors. CE is collisional energy.
